# The *C. elegans* proteome response to two protective *Pseudomonas* mutualists

**DOI:** 10.1101/2023.03.22.533766

**Authors:** Barbara Pees, Lena Peters, Christian Treitz, Inga K. Hamerich, Kohar A. B. Kissoyan, Andreas Tholey, Katja Dierking

## Abstract

The *C. elegans* natural microbiota isolates *Pseudomonas lurida* MYb11 and *Pseudomonas fluorescens* MYb115 protect the host against pathogens through distinct mechanisms. While *P. lurida* produces an antimicrobial compound and directly inhibits pathogen growth, *P. fluorescens* MYb115 protects the host without affecting pathogen growth. It is unknown how these two protective microbes affect host biological processes. We used a proteomics approach to elucidate the *C. elegans* response to MYb11 and MYb115. We found that both *Pseudomonas* isolates increase vitellogenin protein production in young adults, which confirms previous findings on the effect of microbiota on *C. elegans* reproductive timing. Moreover, the *C. elegans* responses to MYb11 and MYb115 exhibit common signatures with the response to other vitamin B_12_-producing bacteria, emphasizing the importance of vitamin B_12_ in *C. elegans*-microbe metabolic interactions. We further analyzed signatures in the *C. elegans* response specific to MYb11 or MYb115. We provide evidence for distinct modification in lipid metabolism by both mutualistic microbes. We could identify activation of host pathogen defense responses as MYb11-specific proteome signature and provide evidence that the intermediate filament protein IFB-2 is required for MYb115-mediated protection. These results indicate that MYb11 not only produces an antimicrobial compound, but also activates host antimicrobial defenses, which together might increase resistance to infection. In contrast, MYb115 affects host processes such as lipid metabolism and cytoskeleton dynamics, which might increase host tolerance to infection. Overall, this study pinpoints proteins of interest that form the basis for additional exploration into the mechanisms underlying *C. elegans* microbiota-mediated protection from pathogen infection and other microbiota-mediated traits.

## Introduction

In line with the growing general interest in host-microbiota interactions, *Caenorhabditis elegans* has emerged as a model host to study the effect of different food and microbiota bacteria on host metabolism and physiology. The bacteria used in these studies include bacteria that likely are associated with nematodes in their habitat, such as *Comamonas aquatica*, *Bacillus subtilis*, and different *Escherichia coli* strains (reviewed in (1)), and probiotic bacteria of human origin such as *Lactobacillus* and *Bifidobacterium* (reviewed in (2)). The characterization of the *C. elegans* natural microbiome (3, 4) and the creation of the simplified natural nematode microbiota mock community CeMbio (5) initiated a steadily increasing number of recent studies on naturally associated microbes and their interaction with the nematode (reviewed in (4, 6)). While we still know relatively little about the function of the *C. elegans* natural microbiota, several studies highlight an important role of the microbiota in supporting the nematode immune response (e.g. (3, 7–9)).

We previously identified two *Pseudomonas* isolates of the natural *C. elegans* microbiota, which protect the worm from infection with *Bacillus thuringiensis* (*Bt*) through different mechanisms: While *P. lurida* MYb11 produces the antimicrobial secondary metabolite massetolide E and directly inhibits pathogen growth, *P. fluorescens* MYb115 does not seem to directly inhibit pathogen growth and may thus protect the host by indirect, host-dependent mechanisms (9). The contribution of the host response to MYb11- and MYb115-mediated protection is unclear.

*C. elegans* responses to different food bacteria and natural microbiota isolates have been investigated mainly by transcriptome analyses (e.g. (10–14)) and only a few proteome analyses (15, 16). Here, we analyzed the direct effects of the protective Pseudomonads MYb11 and MYb115 on the *C. elegans* proteome. To this end, we employed quantitative proteomics and analyzed both, commonalities and differences, in the *C. elegans* proteomic response to MYb11 and MYb115 and did comparative analyses to previously published microbiota- and pathogen-driven host responses. We validated some of the findings using reporter genes, or mutant analyses and thus pinpointed specific proteins that form the groundwork for deeper research into the different molecular mechanisms that underlie *C. elegans*-microbiota interactions particularly in the context of microbiota-mediated protection against pathogens.

## Materials & Methods

### Strains, maintenance, and preparations

Wild type *C. elegans* N2 and all used *C. elegans* mutants/transgenics, as well as bacteria control *Escherichia coli* OP50, were received from sources indicated in Table S1 and maintained according to standard procedures (17). For each experiment, worms were synchronized by bleaching gravid hermaphrodites with alkaline hypochlorite solution and incubating the eggs in M9 overnight on a shaker.

Spore solutions of pathogenic *Bacillus thuringiensis* strains MYBt18247 (Bt247) and MYBt18679 (Bt679) were prepared following a previously established protocol (18), and stored at -20 °C. Single aliquots were freshly thawed for each inoculation.

*Pseudomonas lurida* MYb11 and *Pseudomonas fluorescens* MYb115 belong to the natural microbiota of *C. elegans* (3) and were stored in glycerol stocks at -80 °C. Before each experiment, bacterial isolates were streaked from glycerol stocks onto TSB (tryptic soy broth) agar plates, grown for 2 days at 25 °C, and consequently for an overnight in TSB at 28 °C in a shaking incubator. One day before adding the worms, bacteria of the overnight cultures were harvested by centrifugation, resuspended in 1x PBS (phosphate-buffered saline), pH7, adjusted to an OD_600_ of 10, and used for inoculation of peptone-free medium (PFM, nematode growth medium without peptone) plates.

### qRT-PCR

Worms were raised on OP50, MYb11, or MYb115 at 20 °C until they reached young adulthood, 70 h after synchronized L1s were transferred to the plates. For each replicate roughly 1,000 worms were washed off the plates with 0.025% Triton X-100 in M9 buffer along with three gravity washing steps. Freezing and RNA isolation was done following the instructions of the NucleoSpin RNA/Protein Kit (Macherey-Nagel; Düren, Germany). 1 µg of the extracted total RNA per sample was reverse transcribed using oligo(dt)18 primers (First Strand cDNA Synthesis Kit; ThermoFisher Scientific; Waltham, USA), and 1 µL cDNA was used for qPCR with *tgb-1* as housekeeping gene (19). The expression levels of all tested primers were determined using the iQ™ SYBR® Green Supermix (Bio-Rad, Hercules, USA) using the settings as suggested in the manual. Primer sequences are found in Table S2. The 2^-ΔΔCt^ method was used to calculate the relative gene expression (20).

### Survival and lifespan experiments

For survival experiments, synchronized L1 larvae were grown on PFM plates prepared with lawns of OP50, MYb11, or MYb115 at 20 °C as described above. PFM infection plates were inoculated with serial dilutions of *Bt* spores mixed with bacterial OP50, MYb11, or MYb115 solutions. As L4s, worms were rinsed off the plates and washed with M9 and pipetted in populations of approximately 30 worms on each *Bt* infection plate. After 24 h incubation at 20 °C survival of worms was scored. Worms were considered to be alive when they moved upon gentle prodding with a worm pick. Replicates with less than 15 worms at the time of scoring were excluded.

For lifespan experiments, synchronized L4 larvae were picked onto NGM plates seeded with OP50. Worm survival was determined every day and the alive adults were transferred to new NGM plates with OP50 until the end of the egg laying period.

### Worm imaging and quantification

For imaging of *in vivo* gene/protein expression, transgenic worms were treated as for survival experiments but without *Bt* infection. Young adults (24 h post L4) were then anesthetized with 10 mM tetramisole, placed onto slides containing a fresh 2 % agarose patch, and imaged with a Leica stereomicroscope M205 FA (Wetzlar, Germany). Magnification and exposure time for the fluorophore signal were kept the same in each experiment to ensure comparability; contrast and brightness were adjusted for representative images (grouped worms).

Gene expression of reporter strains was quantified using ImageJ v1.53t (21). Young adults (24 h post L4) were individually imaged and the integrated density (IntDen) of each worm was measured. To correct for potential worm size differences IntDen values were normalized by the total area of each respective individual.

### Proteome analysis

Worms for proteomic analyses were grown on PFM plates prepared with lawns of OP50, MYb11, or MYb115 at 20 °C as described above. L4 stage larvae were transferred to freshly inoculated PFM plates to provide sufficient food. Approximately 1,500 worms per replicate were harvested at 12 h post L4 and washed across a Steriflip® 20 μm nylon mesh filter (Merck; Darmstadt, Germany) with M9 buffer. The samples were prepared as four independent biological replicates.

To each sample, 200 μL protein lysis buffer (100 mM triethylammonium bicarbonate TEAB, 2% SDS, 5 M guanidinium chloride, 2 mM dithiothreitol DTT; 2x complete protease inhibitor) and approximately 200 μL of acid washed glass beads were added. The samples were homogenized using a Bioruptor pico for 20 cycles of 30 s sonication and 30 s cooling at 4 °C. The protein concentration was determined by BCA assay. The proteins were reduced with 10 mM DTT for 1 h at 60 °C and alkylated with 25 mM chloroacetamide at 20 °C for 20 min. The samples were centrifuged for 10 min at 10,000 g and aliquots of 100 μg were prepared following the SP3 protocol (22).

A detailed description of the LC-MS analysis is provided in Supplemental Materials and Methods. Briefly, for each of the 12 samples, approximately 1 µg of peptides were analyzed by liquid chromatography-electrospray ionization-mass spectrometry (LC-ESI MS/MS). Proteome digests were separated over a 2 h gradient on a 50 cm C18 nano-uHPLC column and high-resolution mass spectra were acquired with an Orbitrap Fusion Lumos mass spectrometer. Proteome Discoverer software and the Sequest algorithm were used for peptide identification and label-free quantification. MS data were searched against the reference proteome of *C. elegans* (26,738 entries) combined with the UniParc entries of *P. lurida* (5,392 entries), *P. fluorescens* (5,548 entries), and *E. coli* OP50 (4,227 entries). Statistical evaluation of the quantitative data was performed with the Perseus software (23). LC-MS raw data were deposited to the ProteomeXchange Consortium via the PRIDE partner repository (24) with the dataset identifier PXD040520.

### Statistical analyses

For identification of differentially abundant proteins, we performed a one-way ANOVA comparing the three conditions (OP50 *vs* MYb11 *vs* MYb115) and corrected for multiple comparisons using a permutation-based FDR analysis. An FDR cut-off of 5% was applied and Tukey’s HSD test was used for *post hoc* analysis. Significant protein groups assigned to each of the pairs of conditions were tested for UniProt keywords by Fisher exact test corrected for multiple testing by Benjamini-Hochberg FDR calculation. All significant findings with an FDR below 5% are provided in Table S3.

Heatmaps were created using the Morpheus (https://software.broadinstitute.org/morpheus), GO term overrepresentation analyses were done with eVitta v1.3.1 (25). All remaining statistical analyses were carried out with RStudio, R v4.2.1, graphs created with its package ggplot2 v3.3.6 (26), and edited with Inkscape v1.1.2.

## Results

### Common proteomic response to protective *Pseudomonas*

We were interested in identifying the proteomic changes in *C. elegans* exposed to two protective *Pseudomonas* isolates, *P. lurida* MYb11 and *P. fluorescens* MYb115. To this end, worms were grown on MYb11, MYb115, or *E. coli* OP50 and harvested for proteome analysis as young adults. Using LC-MS analysis we identified 4,314 protein groups in total, which included 259 protein groups annotated to bacterial taxa and 4,055 to *C. elegans* protein groups. For statistical evaluation, the identified *C. elegans* proteins were filtered to 3,456 entries quantified in all four replicates of at least one bacterial treatment. The complete list of proteins is provided in Table S3.

Comparing MYb-treated worms to those grown on OP50, 674 proteins were differentially abundant. Among these proteins, 201 showed a significant difference in both *Pseudomonas* treatments, MYb11 *vs* OP50 and MYb115 *vs* OP50 (Figure 1A; Table S3). When we grouped the shared proteomic response towards *Pseudomonas* into more and less abundant proteins, we obtained 84 higher abundant proteins and 104 less abundant proteins (Figure 1B, C). Strikingly, among the more abundant proteins we found all six vitellogenins described in *C. elegans* (27, 28). Vitellogenins are yolk proteins which are primarily produced in the reproductive phase to supply energy to the embryos (29). Expression of the vitellogenins encoding *vit* genes is known to be greatly up-regulated in young adults and down-regulated in aging worms (30). We have previously shown that MYb11 and MYb115 accelerate *C. elegans* reproductive maturity without affecting the overall reproductive output (31). Thus, it might be possible that the abundance of vitellogenins in worms treated with either of the Pseudomonads reflects these differences in reproductive maturity. When we compared the abundance of the vitellogenin VIT-2 between young adults on MYb11, MYb115, or OP50 using a *C. elegans vit-2*::gfp reporter strain, we indeed observed an increased number of VIT-2 expressing eggs/embryos and VIT-2 abundance in worms on MYb11 and MYb115 (Figure 1B). This observation is reminiscent of data on *Comamonas aquatica* DA1877 and *Enterobacter cloacae* CEent1 that accelerate *C. elegans* development (7, 10).

**Figure 1.**
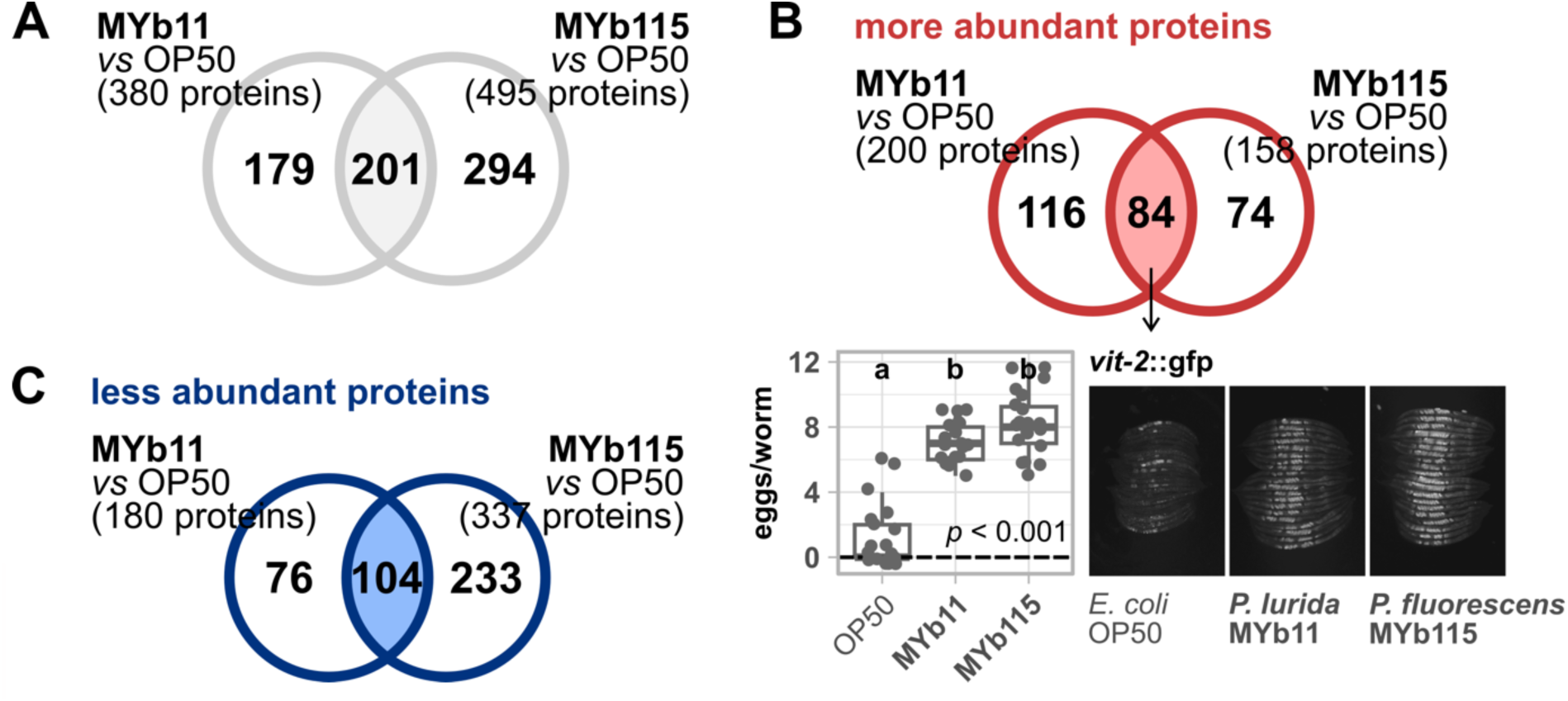
Proteomic response of *C. elegans* towards mutualistic *Pseudomonas*. Venn diagrams showing (A) all significantly differentially abundant proteins resulting from comparing either MYb11-exposed worms to OP50-exposed worms or MYb115-exposed worms to OP50-exposed worm, (B) only the significantly more abundant proteins, or (C) the significantly less abundant proteins; ANOVA, post hoc Tukey HSD, p > 0.05. (B) Transgenic *C. elegans* reporter strain *vit-2*::gfp demonstrating *in vivo* abundance of VIT-2. Worms were exposed to either *E. coli* OP50, *P. lurida* MYb11, or *P. fluorescens* MYb115, and gfp signals imaged in groups of 20 individuals as young adults. Worms were arranged with the heads pointing to the right. The boxplot displays the quantification of VIT-2 expressing eggs/embryos in young adults (24 h post L4). Each dot represents one worm with n = 20, the dashed line represents the median number of eggs per worm for OP50-exposed worms. The *p*-value indicates the statistical significance among the differently exposed worms according to a Kruskal-Wallis rank sum test (32). The *post hoc* Dunn’s test (33) with Bonferroni correction provides the statistical significances between the differently exposed worms and is denoted with letters (same letters indicate no significant differences). Raw data and corresponding *p*-values are provided in Table S6.

### Microbiota bacteria elicit a robust proteomic response related to vitamin B_12_- dependent metabolism

*Pseudomonas* and *Ochrobactrum* represent the most prevalent genera in the natural *C. elegans* microbiota, are able to colonize the host, and seem to have largely beneficial effects on host life-history traits (3–5, 9, 34). We previously analyzed the effects of *O. vermis* MYb71 and *O. pseudogrignonense* MYb237 on the *C. elegans* proteome (15). Here, we asked whether the *C. elegans* proteome response to the Pseudomonads MYb11 and MYb115 shares common signatures with the response to *O. vermis* MYb71 and *O. pseudogrignonense* MYb237. We extracted significantly differentially abundant proteins in either MYb71 *vs E. coli* OP50 or MYb237 *vs E. coli* OP50 from the published dataset, and examined the overlap between responses to all four microbiota isolates. We identified 32 proteins, whose abundances were affected by all four microbiota bacteria (Figure 2A; Table S6). 31 of the 32 proteins showed a common increase and decrease in abundances, respectively, relative to the control *E. coli* OP50. One protein, the uncharacterized CHK domain-containing protein F58B4.5, was more abundant in worms fed with either *Ochrobactrum* isolates but less abundant on *Pseudomonas*. It thus represents a promising candidate for understanding contrasting responses to both taxa. We further noticed that 11 proteins out of the 31 proteins representing the common proteome response to *Pseudomonas* and *Ochrobactrum* are members of the interacting methionine/S-adenosylmethionine (met/SAM) cycle, which is part of the one-carbon cycle, and the alternative propionate shunt pathway (35, 36) (Figure 2B). In this signaling network, vitamin B_12_ is a crucial micronutrient that feeds into methionine synthesis and allows the breakdown of propionate (35, 37), thereby promoting *C. elegans* longevity, fertility, development, and mitochondrial health (38, 39).

**Figure 2.**
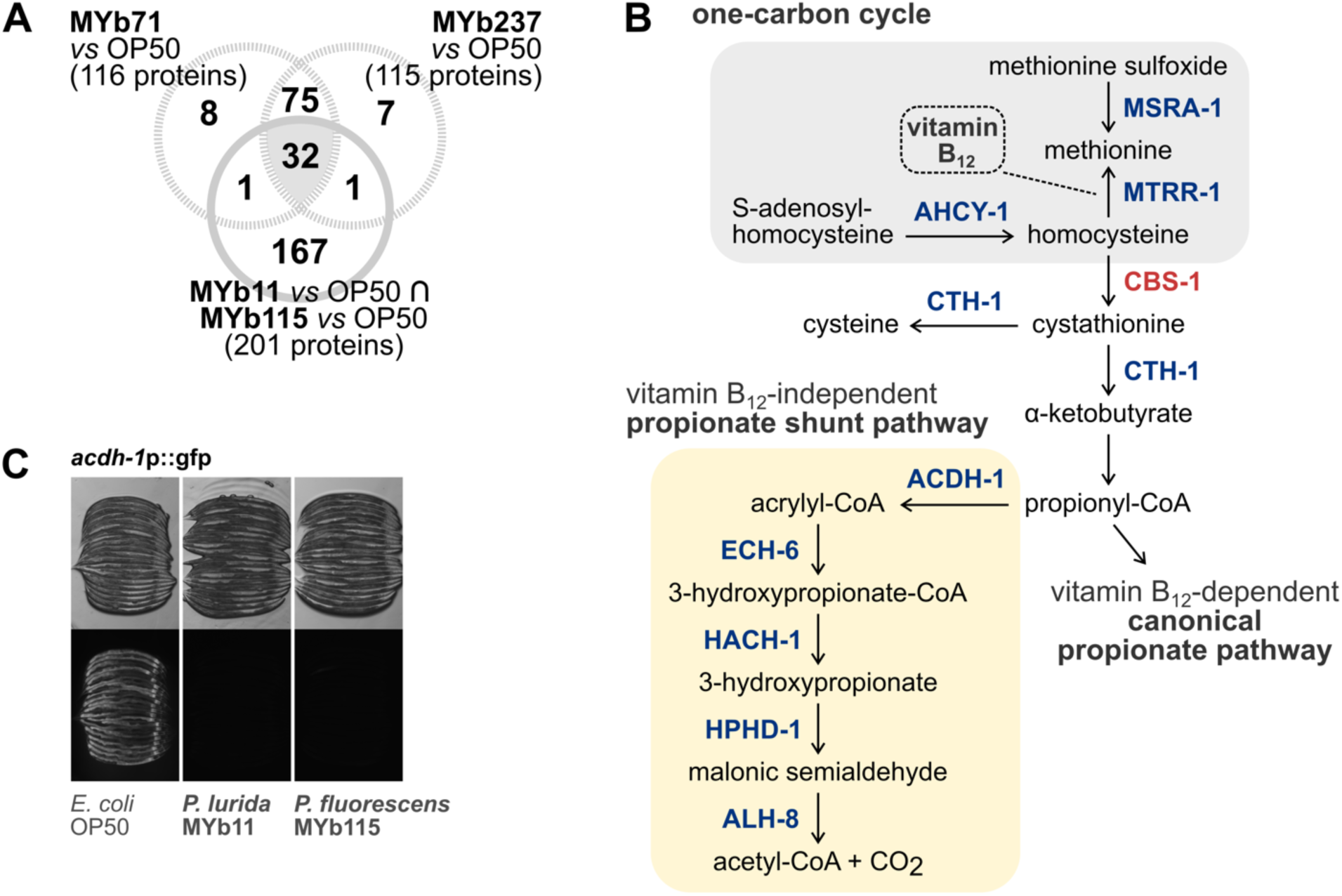
Changes in vitamin B12-dependent metabolism are shared proteomic responses to mutualistic *Pseudomonas* and *Ochrobactrum.* (A) Venn diagram showing all significantly differentially abundant proteins resulting from the overlap of the comparison MYb11 *vs* OP50 and MYb115 *vs* OP50 (Figure 1A) compared against differentially abundant proteins on *Ochrobactrum* MYb71 and MYb237. (B) Excerpt of the one-carbon cycle (gray background) and the propionate pathways (yellow background). Shown are mainly the steps which involve commonly differentially abundant proteins in worms grown on mutualistic *Pseudomonas* and *Ochrobactrum*. Protein coloring depicts either less abundant (blue) or more abundant (red) proteins. CoA = coenzyme A. Adapted from (35, 36). (C) Transgenic *C. elegans* reporter strain *acdh-1*p::gfp demonstrating *in vivo* expression of *acdh-1*. Worms were exposed to either *E. coli* OP50, *P. lurida* MYb11, or *P. fluorescens* MYb115, and gfp signals imaged in groups of 20 individuals as young adults. Worms were arranged with the heads pointing to the right; transmission light images in upper panel corresponding to fluorescence images in lower panel.

In the presence of vitamin B_12_ *C. elegans* uses the canonical propionate pathway to degrade propionate into less toxic metabolites and, simultaneously, inactivates the B_12_-independent propionate shunt, i.e. by down-regulating the partaking genes (35, 40) (Figure 2B). Exactly these propionate shunt proteins, ACDH-1, ECH-6, HACH-1, HPHD-1, and ALH-8, were less abundant in the microbiota-treated worms which is evidence for the provision of vitamin B_12_ by *Pseudomonas* and *Ochrobactrum*. Also, genes encoding the 12 proteins that show different abundances by *Pseudomonas* and *Ochrobactrum* (Table S6), were reported to be differentially regulated by either *C. aquatica* DA1877 or vitamin B_12_ supplementation (35, 36, 41). We confirmed that expression of the acyl-CoA dehydrogenase encoding gene *acdh-1* is down-regulated by MYb11 and MYb115 by using the dietary sensor *C. elegans* strain *acdh-1*p::gfp, which reacts to vitamin B_12_ presence (35, 42) (Figure 2C).

### Proteomic responses of *C. elegans* specific to MYb11 and MYb115

While both Pseudomonads, MYb11 and MYb115, are able to protect *C. elegans* from *Bt* infection, the underlying mechanisms are distinct (9). As a step toward understanding the contribution of the host response to MYb11- and MYb115-mediated protection, we sought to identify the differences in the proteomic responses between worms exposed to MYb11 and MYb115. Both treatments were directly compared and we found 421 proteins that differed significantly in their abundance between the two conditions (Figure 3A). Interestingly, 326 proteins were more abundant in worms grown on MYb11 compared to MYb115 and only 95 proteins were more abundant in MYb115-exposed worms compared to MYb11-exposed worms (Figure 3A). To extract the proteins that were uniquely differently abundant in either MYb11 or MYb115, we included the data on OP50 to generate 4 clusters using *k*-means clustering: cluster 1 and 4 represent proteins whose abundance only changed in MYb11-exposed worms, i.e., in reference to MYb115 and OP50, while cluster 2 and 3 represent proteins with different abundances specifically in MYb115-exposed worms, i.e., in reference to MYb11 and OP50 (Figure 3A). Next, we employed eVitta, an online tool developed for the analysis and visualization of transcriptome data (25), to look for enriched gene ontology (GO) terms in these clusters.

**Figure 3.**
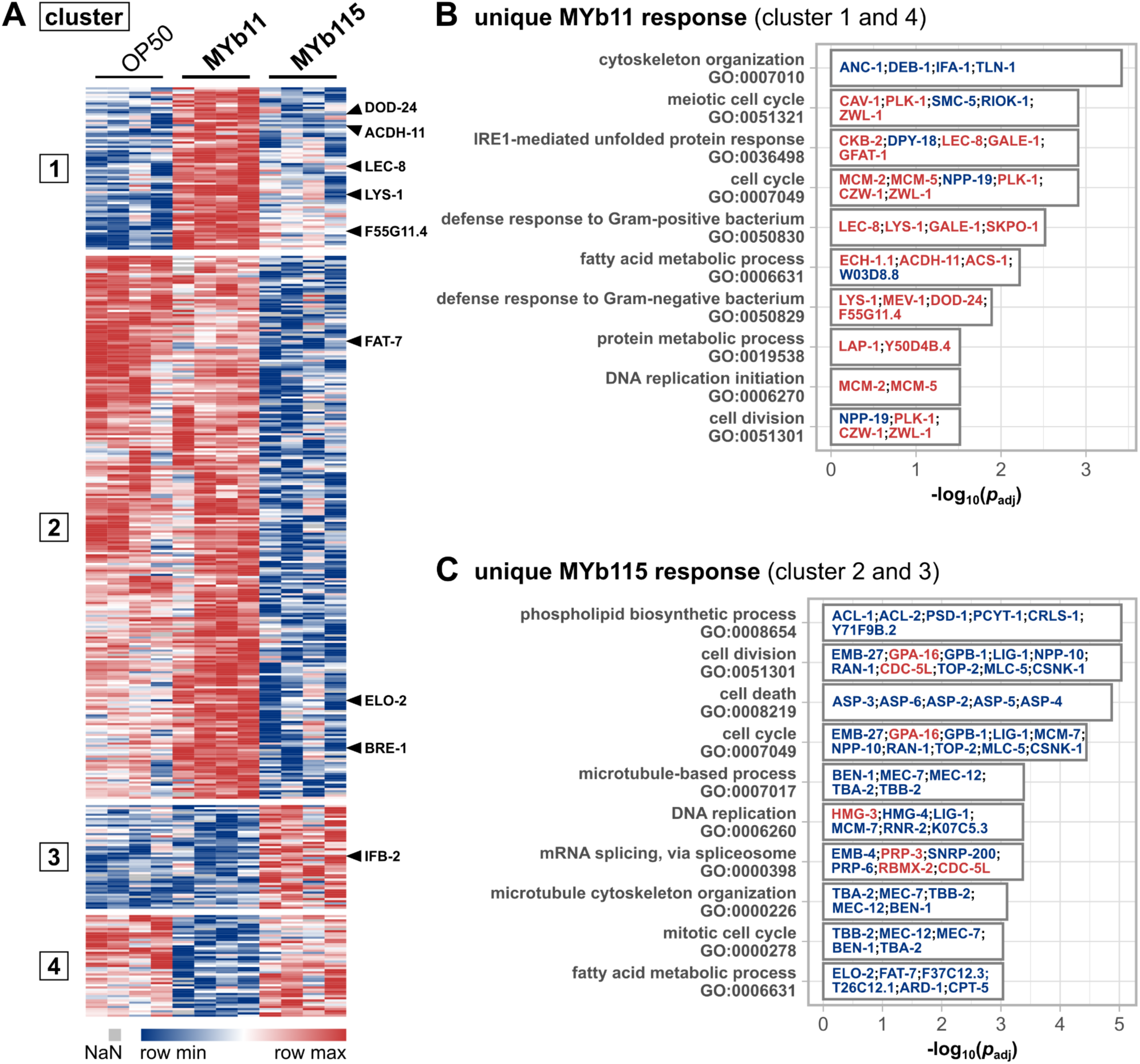
Differences in the proteomic responses of *C. elegans* towards *P. lurida* MYb11 compared to *P. fluorescens* MYb115. (A) Heatmap showing the log2 label-free intensity values of differentially abundant proteins in the comparison of MYb11-exposed worms against MYb115-exposed worms. The columns denote the bacterial treatment with 4 replicates each, each row represents one protein. By including the data on OP50, abundance values were separated into 4 clusters using the *k*-means clustering approach. The rows of exemplary proteins mentioned in the text are marked on the heatmap’s right. Bar plot of significantly enriched gene ontology (GO) terms in either (B) clusters 1 and 4 (different abundances of proteins uniquely in MYb11-treated worms) or (C) clusters 2 and 3 (different abundances of proteins uniquely in MYb115-treated worms). The proteins which are assigned to the respective GO term are noted on the bars, their coloring indicates higher (red) or lower (blue) abundance. Shown are the 10 GO terms with the highest significance. The complete list of GO terms is to be found in Tables S4 and S5.

### MYb11 causes a mild pathogen response in *C. elegans*

Proteins affected by both *Pseudomonas* isolates were enriched in GO terms associated with nucleic acids (e.g., DNA replication, mRNA splicing), and also fatty acid-related terms albeit targeting different fat metabolism enzymes (further discussed in next paragraph). On the contrary, defense response proteins were a MYb11-linked feature with defense responses to Gram-positive bacterium (GO:0050830) and Gram-negative bacterium (GO:0050829) among the 10 highest significantly enriched GO terms in the unique MYb11 response (Figure 3B; Table S4). Interestingly, the 7 proteins (LYS-1, LEC-8, GALE-1, SKPO-1, MEV-1, DOD-24, F55G11.4) associated with the GO defense response terms were all more abundant in MYb11 compared to MYb115 (Figure 3C; Table S5), indicating that MYb11 induces *C. elegans* pathogen defenses while MYb115 does not. This finding is in line with the previous observation that MYb11 has a pathogenic potential in some contexts, despite its protective effect against *Bt* and *P. aeruginosa*, resulting in a shorter lifespan and increased susceptibility to purified *Bt* toxins (31). Interestingly, the lifespan of MYb11-exposed worms on nutritious medium (NGM) (Figure S1) is much more decreased than on minimal medium (PFM), suggesting that the detrimental effect on worms is primarily promoted by proliferating and metabolically active MYb11. Hence, we assessed the general pathogenic potential of MYb11 and MYb115 and tested activation of the *C. elegans* stress reporters, *hsp-4*::gfp (endoplasmic reticulum stress), *hsp-6*::gfp and *hsp-60*::gfp (mitochondrial stress (43, 44)), *gst-4*p::gfp (oxidative stress (45)), and the immune reporters *irg-1*p::gfp (46) and *clec-60*p::gfp (47) (Figure S2). Bacteria from the natural *C. elegans* habitat were reported to induce expression of some of these reporter genes (48). We found that the oxidative stress reporter *gst-4*p::gfp was significantly up-regulated only by MYb11 (Figure S2). MYb11 also slightly induced expression of the C-type lectin-like gene *clec-60* reporter compared to OP50, however, only significantly when compared to MYb115-mediated induction. These results indicate that mainly MYb11 activates the *C. elegans* oxidative stress response and the expression of *clec-60*p::gfp. To explore in how far the *C. elegans* induced proteome response to MYb11 overlaps with the induced proteome response to pathogenic bacteria, we compared our data (Figure 1B) with the proteomic changes elicited by pathogenic *P. aeruginosa* PA14 (49) and Bt247 (50). The comparison of proteins of higher abundance in MYb11-exposed worms with PA14-responsive proteins yielded an overlap of 14 more abundant proteins (Figure 4A). Among these 14 proteins were the known pathogen-responsive CUB-like domain proteins C17H12.8, C32H11.4, DOD-17, DOD-24 and F55G11.4, and the infection response gene (IRG) 3. Similarly, when we compared the response to MYb11 to the proteomic response to *Bt* infection, the abundances of 11 proteins were commonly increased (Figure 4B). Among these proteins were the CUB-like domain proteins C17H12.8, the lysozyme LYS-1, the galectins LEC-8 and LEC-9, and the C-type lectin-like domain proteins CLEC-41 and CLEC-65. Notably, most of these MYb11- and pathogen-responsive proteins were indeed less responsive to MYb115 (Figure 4A, B).

**Figure 4.**
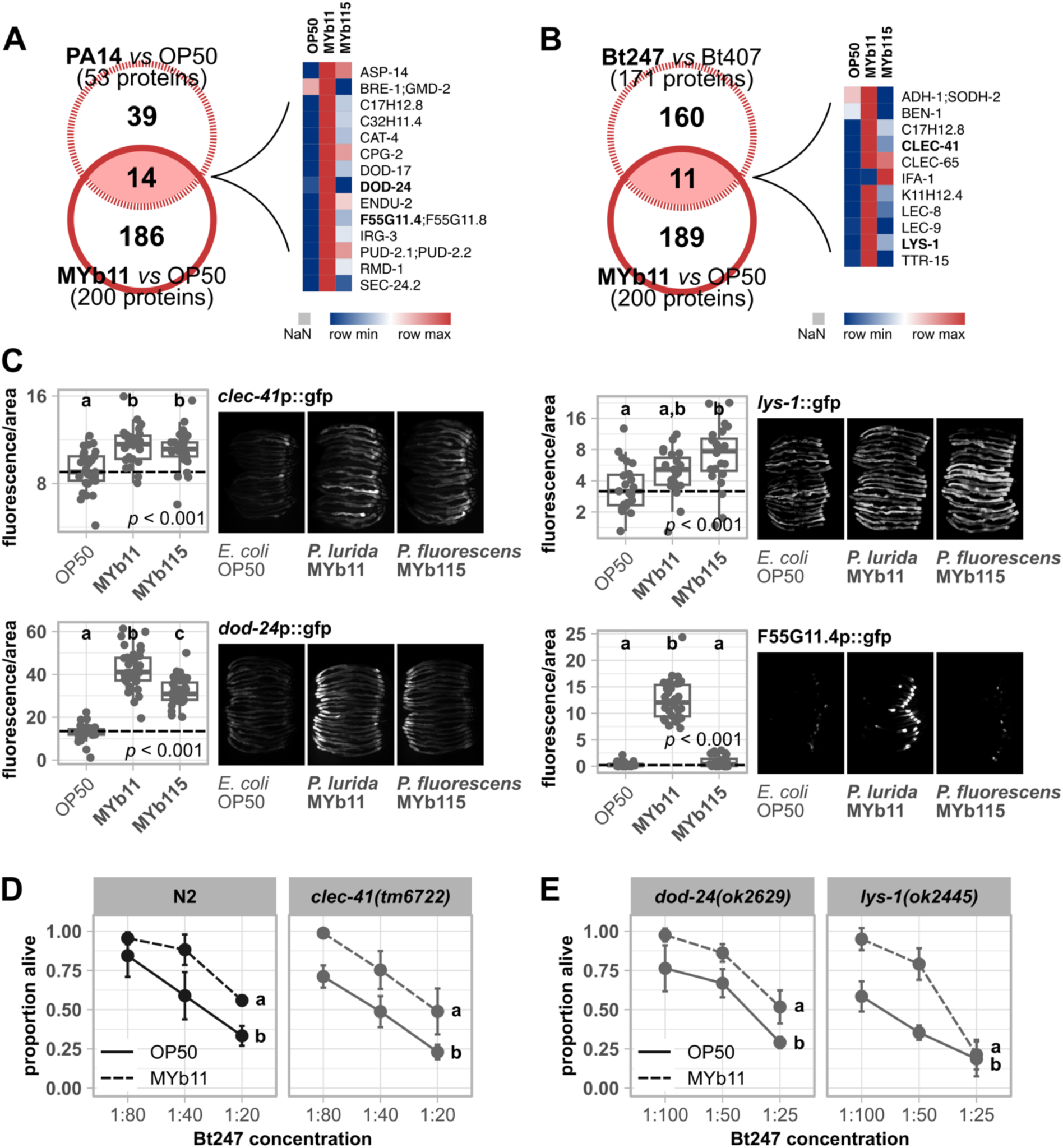
MYb11 activates expression of *C. elegans* innate immune response genes and proteins. Venn diagrams showing significantly more abundant proteins resulting from the comparison MYb11 *vs* OP50 (Figure 1B) compared against significantly more abundant proteins of (A) *P. aeruginosa* PA14 *vs E. coli* OP50, and (B) *B. thuringiensis* Bt247 *vs* non-pathogenic strain Bt407. The accompanying heatmaps represent the averaged log2 label-free intensity values (*n* = 4) of the overlapping significant proteins. Data taken from (49, 50). (C) Transgenic *C. elegans* reporter strains demonstrating *in vivo* expression of selected promotor sequences tagged with gfp. Transgenic strains were exposed to either *E. coli* OP50, *P. lurida* MYb11, or *P. fluorescens* MYb115, and fluorescent signals imaged in groups of 20 individuals as young adults. Worms were arranged with the heads pointing to the right. The boxplots display the quantification of the gfp fluorescence in young adults (24 h post L4) normalized by the worm’s body size (area). Each dot represents one worm with *n* = 29-35, the dashed line represents the median of the mean grey value for OP50-exposed worms. The *p*-value indicates the statistical significance among the differently exposed worms according to a Kruskal-Wallis rank sum test (32). The *post hoc* Dunn’s test (33) with Bonferroni correction provides the statistical significances between the differently exposed worms and is denoted with letters (same letters indicate no significant differences. (D, E) Survival of mutants *clec-41(tm6722)*, *dod-24(ok262*9), and *lys-1(ok2445)* and wild type N2 infected with serial dilutions of *B. thuringiensis* Bt247 after 24 hpi (post infection). Worms were exposed to either OP50 or MYb11, before and during infection. Each dot represents the mean ± standard deviation (SD) of four worm populations (*n* = 4). Same letters indicate no significant differences between the dose response curves according to a generalized linear model (GLM) (51) and Bonferroni correction. Raw data and corresponding *p*-values are provided in Table S6, an additional repetition of the experiment (D) is to be found in Figure S4.

Although MYb11 produces the antimicrobial secondary metabolite massetolide E and directly inhibits pathogen growth (9), activation of host-pathogen defense responses, i.e. production of host immune proteins, may contribute to MYb11-mediated protection. To explore this possibility, we focused on F55G11.4, DOD-24, LYS-1, and CLEC-41, whose abundances were strongly increased by MYb11. F55G11.4 was the protein with the highest abundance on MYb11 (Table S3). DOD-24 is commonly used as marker of the immune response to PA14 and other Gram-negative pathogens (7, 52–54). LYS-1 is required for normal resistance to the Gram-positive *Staphylococcus aureus* (55), and CLEC-41 has demonstrated immune effector function and exhibits antimicrobial activity against Bt247 *in vitro* (56). Mutants of all genes, but F55G11.4, were available at the CGC. First, using qRT-PCR and gfp reporter gene promoters, we confirmed that expression of *dod-24* and F55G11.4 is significantly up-regulated by MYb11 in comparison to MYb115 or OP50 also on the transcript level (Figure 4C; Figure S3). The expression of the *lys-1* reporter, however, was increased by both MYb11 and MYb115, albeit significantly only by MYb115, and expression of the *clec-41* reporter was significantly induced by both Pseudomonads (Figure 4C). To determine if these MYb11-induced genes have a function in MYb11-mediated protection against *Bt* infection we grew the available *dod-24, clec-41, and lys-1* knock-out mutants on OP50, MYb11, or MYb115, infected them with Bt247, and scored their survival. MYb11 increased resistance to Bt247 infection also in *dod-24, clec-41, and lys-1* mutants (Figure 4D, E; Figure S4).

### MYb11 and MYb115 cause diverging responses in *C. elegans* fat metabolism

Among the 10 highest significantly enriched GO terms concerning biological processes in the unique MYb11 response as well as in the unique MYb115 response, we found the term fatty acid metabolic process (GO:0006631) (Figure 3B, C). Moreover, the GO term phospholipid biosynthetic process (GO:0008654) was enriched only in the unique MYb115 response (Figure 3C). Since the ability to mount an immune response has been repeatedly linked to changes in *C. elegans* fat metabolism (e.g. (57, 58)), we took a closer look at the underlying proteins. While the predicted fatty acid β-oxidation enzyme ECH-1.1, the acyl-CoA dehydratase ACDH-11, and the acyl-CoA synthetase ACS-1 were of higher abundance in worms on MYb11 (Figure 3B; Table S3), the fatty acid elongase ELO-2 and the fatty acid desaturase FAT-7 were of lower abundance in worms on MYb115 (Figure 3C; Table S3). FAT-6 and FAT-7 are members of the long-chain fatty acid synthesis pathway and act redundantly in the synthesis of the monounsaturated fatty acid oleate from stearic acid (59). We validated the effect of the *Pseudomonas* isolates on FAT-7 by assessing the *in vivo* protein abundance of *fat-7*::gfp in worms exposed to *E. coli* OP50, *P. lurida* MYb11, or *P. fluorescens* MYb115. Expression of *fat-7*::gfp was indeed significantly reduced in worms on MYb115 compared to worms on OP50 or MYb11 (Figure 5A), confirming that MYb11 and MYb115 cause diverging responses in *C. elegans* fat metabolism.

**Figure 5.**
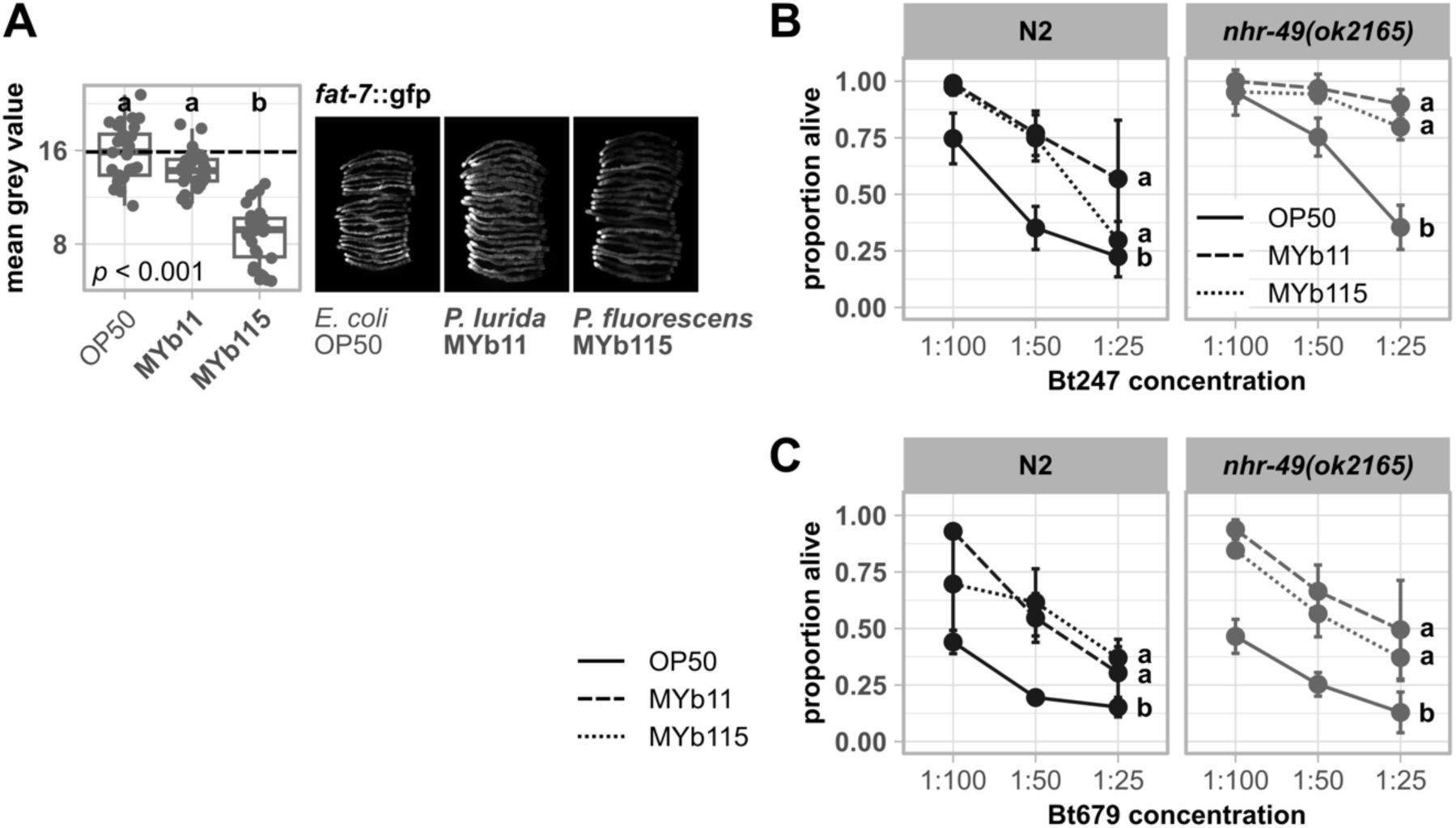
Divergent proteomic changes in fat metabolism occur in MYb11- and MYb115-exposed worms but common fat metabolism regulator NHR-49 is not involved in the defense against *Bt* infection. (A) Transgenic *C. elegans* reporter strain demonstrating *in vivo* abundance of FAT-7. Worms were exposed to either *E. coli* OP50, *P. lurida* MYb11, or *P. fluorescens* MYb115, and gfp signals imaged in groups of 20 individuals as young adults. Worms were arranged with the heads pointing to the right. The boxplots display the quantification of the gfp fluorescence in young adults (24 h post L4) normalized by the worm’s body size (area). Each dot represents one worm with *n* = 29-30, the dashed line represents the median of the mean grey value for OP50-exposed worms. The *p*-value indicates the statistical significance among the differently exposed worms according to a Kruskal-Wallis rank sum test (32). The *post hoc* Dunn’s test (33) with Bonferroni correction provides the statistical significances between the differently exposed worms and is denoted with letters (same letters indicate no significant differences). (B) Survival of mutant *nhr-49(ok2165)* and wild type N2 infected with serial dilutions of (B) *B. thuringiensis* Bt247 or (C) Bt679 after 24 hpi. Worms were fed with either OP50, MYb11, or MYb115 before and during infection. Each dot represents the mean ± standard deviation (SD) of (B) four or (C) three worm populations (*n* = 3-4). Same letters indicate no significant differences between the dose response curves according to a generalized linear model (GLM) (51) and Bonferroni correction. Raw data and corresponding *p*-values are provided in Table S6, additional repetitions of the same experiments are to be found in Figure S5.

The nuclear hormone receptor NHR-49 is a major regulator of *C. elegans* fat metabolism and activates *fat-7* expression (60). Thus, we evaluated the role of *nhr-49* in the protective effect mediated by either *Pseudomonas* isolate. We tested the survival of the knock-out mutant *nhr-49(ok2165)* infected with the *Bt* strain Bt247 or Bt679, in the presence of either OP50, MYb11, or MYb115. Neither MYb11-, nor MYb115-mediated protection against *Bt* infection was dependent on *nhr-49* (Figure 5B, C; Figure S5).

### Intermediate filament IFB-2 may be involved in MYb115-mediated protection against *B. thuringiensis*

Another intriguing result of our overrepresentation analysis was the presence of cytoskeleton-related terms (e.g., GO:0007010, GO:0007071, GO:0000226) (Figure 3B, C). As our previous proteome dataset of *C. elegans* infected with *B. thuringiensis* similarly showed an enrichment in cytoskeleton-based GO terms (50, 61), we wondered whether systematic reorganization of the cytoskeleton evoked by microbiota members MYb11 and MYb115 might mediate defense against *Bt*. Therefore, we extracted all proteins of our proteome dataset with the GO term cytoskeleton (Table S3) and analyzed their abundance pattern (Figure 6A). Strikingly, four out of five intermediate filaments we identified in the overall analysis, IFB-2, IFP-1, and two IFA-1 isoforms, were more abundant in MYb115-treated worms compared to MYb11- or OP50-fed worms.

**Figure 6.**
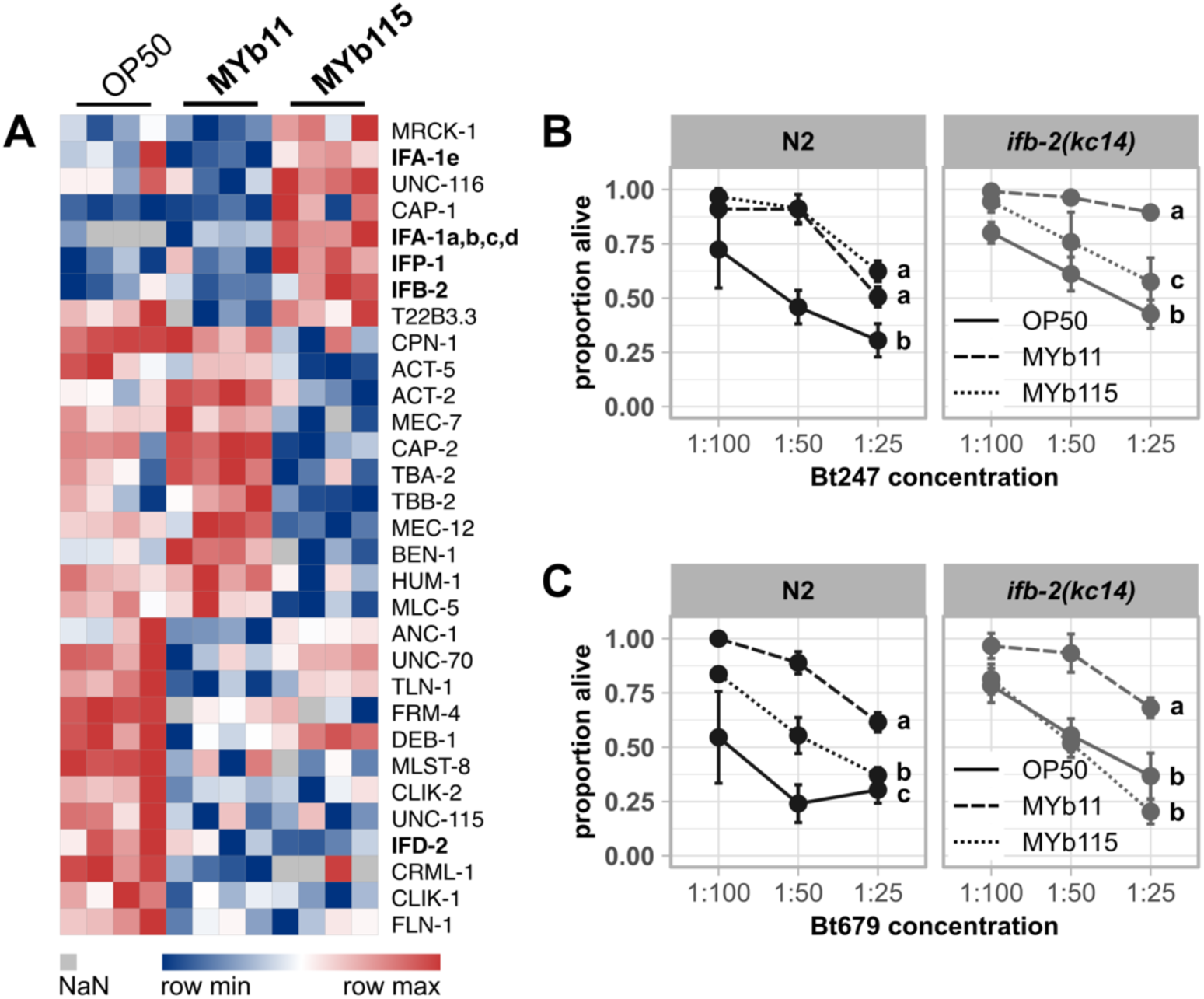
MYb115-mediated protection against *Bt* infection may depend on IFB-2. (A) Heatmap showing the log2 label-free intensity values of identified proteins related to the GO term cytoskeleton. The columns denote the bacterial treatment with 4 replicates each, each row represents one protein. Survival of wild type N2 and mutant *ifb-2(kc14)* infected with serial dilutions of (B) *B. thuringiensis* Bt247 or (C) Bt679 after 24 hpi. Worms were exposed to either OP50, MYb11, or MYb115 before and during infection. Each dot represents the mean ± standard deviation (SD) of (B) four or (C) three worm populations (*n* = 3-4). Same letters indicate no significant differences between the dose response curves according to a generalized linear model (GLM) (51) and Bonferroni correction. Raw data and corresponding *p*-values are provided in Table S6, additional repetitions of the same experiments are to be found in Figure S6.

The cytoskeleton, consisting of actin-based microfilaments, tubulin-based microtubules, and intermediate filaments (62), canonically stabilizes and maintains the cellular shape ((63); reviewed in (64)). The six *C. elegans* intestinal intermediate filaments, IFB-2, IFC-1, IFC-2, IFD-1, IFD-2, and IFP-1 are all located in the endotube (65), which is positioned at the interface between the intestinal brush border and the cytoplasm (66). To determine the contribution of intermediate filament proteins in the endotube to microbiota-mediated protection against Bt247 and Bt679 infection, we tested the *ifb-2(kc14)* mutant, which completely lacks an endotube (66). We found that the protective effect of MYb115 against *Bt* infection is indeed either partially (Figure 6B), or completely abolished in the *ifb-2* mutant in four out of five experiments (Figure 6C; Figure S6A, B, D). On the contrary, the MYb11-mediated protective effect seems to be independent of IFB-2 (Figure 6B, C; Figure S6C, D).

## Discussion

This study represents a proteome analysis of the *C. elegans* response to its microbiota members *P. lurida* MYb11 and *P. fluorescens* MYb115 that were previously shown to protect the host against pathogen infection (9). We compared the proteome response elicited by MYb11 and MYb115 with the proteome response to other naturally associated bacteria, to known *C. elegans* pathogens, and directly to each other, to reveal common and specific signatures. We thus identified candidate proteins (Figure 7) that are the basis for further investigation of the mechanisms that mediate pathogen protection.

**Figure 7.**
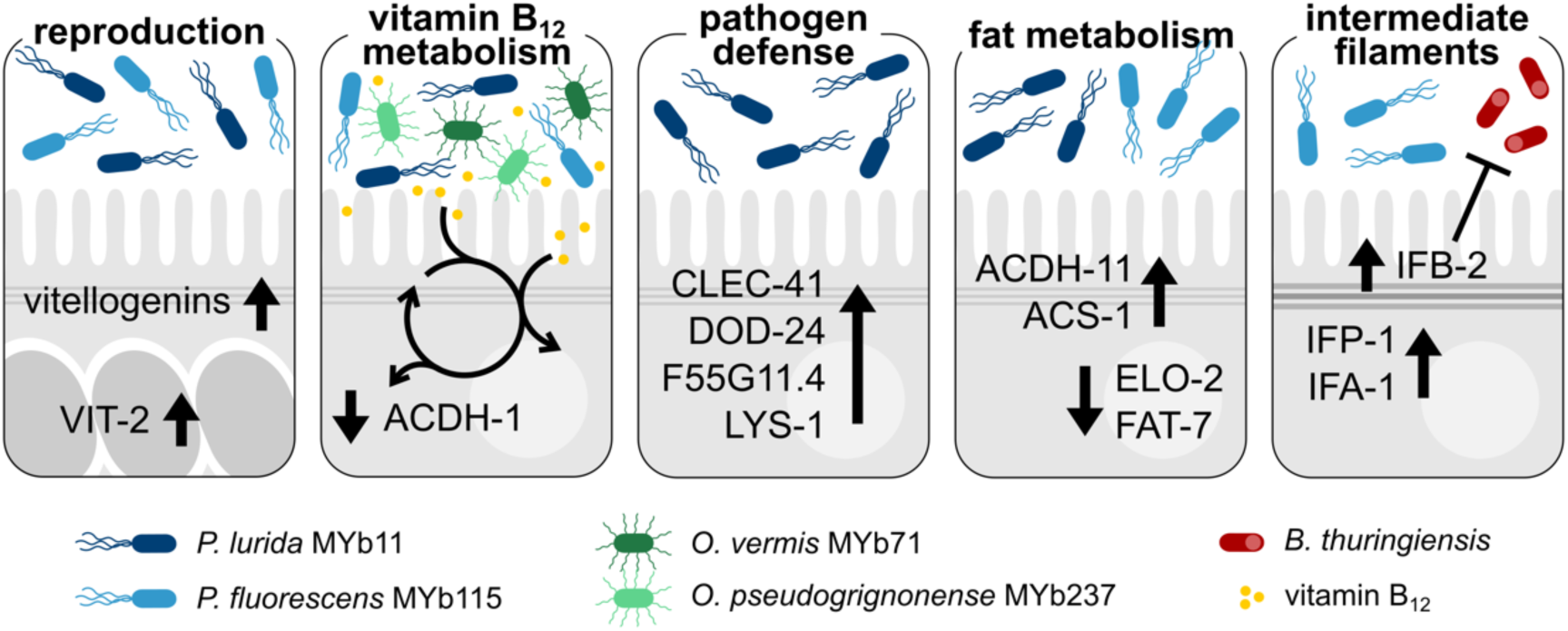
Candidate proteins that are potentially involved in *C. elegans* microbiota-mediated protection. Both *Pseudomonas* isolates, *P. lurida* MYb11 and *P. fluorescens* MYb115, increase *C. elegans* vitellogenin protein production and affect host vitamin B12 metabolism. The latter is also affected by other vitamin B12-producing microbiota bacteria, such as *O. vermis* MYb71 and *O. pseudogrignonense* MYb237. MYb11 activates host-pathogen defense responses more strongly than MYb115. Moreover, both MYb11 and MYb115 modify host fat metabolism, but affect different proteins. MYb115 increases intermediate filament proteins and MYb115-mediated protection against *Bt* infection was reduced in an *ifb-2* mutant.

To reveal common signatures in the *C. elegans* proteome response to naturally associated bacteria, we compared our data with the response to *O. vermis* MYb71 and *O. pseudogrignonense* MYb237, two other members of the *C. elegans* natural microbiota (15). Strikingly, the robust, shared proteomic response of *C. elegans* to mutualistic *Pseudomonas* and *Ochrobactrum* seems to be driven by the availability of vitamin B_12_ and subsequent metabolic signaling: 35% of the commonly affected proteins are members of the interacting met/SAM cycle and the alternative propionate shunt pathway (35, 36). Both *Ochrobactrum* isolates, MYb71 and MYb237, and both *Pseudomonas* isolates, MYb11 and MYb115, are predicted vitamin B_12_ producers (34) and our proteomic analysis corroborates this finding. The importance of microbial-derived vitamin B_12_ in regulating the host met/SAM cycle has previously been demonstrated by comparing a *C. aquatica* DA1877 diet, which is naturally rich in vitamin B_12_, to the standard *C. elegans* laboratory food bacterium *E. coli* OP50 (35). Since *E. coli* OP50, which is low in vitamin B_12_, is also commonly used as control in *C. elegans* microbiota studies, it is important to consider the effect of microbial-derived vitamin B_12_ on *C. elegans* and the resulting, potentially divers effects on host physiology. For example, vitamin B_12_ was identified as the major metabolite accelerating *C. elegans* development and reproductive timing (35, 39). Moreover, vitamin B_12_ can affect regulation of host growth, lifespan, chemosensory receptor gene expression, and responses to stress (10, 38, 67, 68). These and other findings stress the importance of microbe-derived vitamin B_12_ in *C. elegans* metabolic processes, which should be considered when studying the effects of the (potentially vitamin B_12_-producing) *C. elegans* microbiota on host physiology.

We are also interested in placing the *C. elegans* proteome response to MYb11 and MYb115 in the context of microbiota-mediated protection against pathogen infection. Both Pseudomonads protect the worm against *Bt* infection, but in how far the host response contributes to Myb11- and Myb115-mediated protection remains poorly understood (9, 31). Our proteome analyses revealed several interesting host candidate proteins that may be involved in MYb11- and/or MYb115-mediated protection against *Bt*. First, the abundance of all six vitellogenins described in *C. elegans* (27, 28) was affected by both *Pseudomonas* isolates. In addition to their function in energy supply for the developing embryo, vitellogenins may play a role in pathogen defenses. In the honey bee vitellogenin drives transgenerational immune priming by binding pathogen-associated molecular patterns of e.g. *E. coli* and by transporting these signals into developing eggs (69). Also, in *C. elegans*, vitellogenins are involved in defense against *Photorhabdus luminescens* (70). Even more relevant, VIT-2 is required for *Lactobacillus*-mediated protection against Methicillin-resistant *S. aureus*, albeit in aging worms (16). Second, as discussed above, both *Pseudomonas* isolates decrease abundance of proteins of the vitamin B_12_-independent propionate shunt, which indicates that MYb11 and MYb115 provide vitamin B_12_ to the host. Increased vitamin B_12_ availability was shown to improve *C. elegans* mitochondrial health and resistance to infection with *P. aeruginosa* and *Enterococcus faecalis* in a liquid-based killing assay, but not to *P*. *aeruginosa*-mediated slow killing (39). Furthermore, increased vitamin B_12_ availability protects *C. elegans* against exposure to the thiol-reducing agent dithiothreitol (71).

We also identified proteins that were affected by either microbiota isolate. This aspect is of relevance since we know that the protective mechanisms mediated by MYb11 and MYb115 are distinct and that MYb11 and MYb115 have distinct effects on host physiology: MYb11 produces the antimicrobial compound massetolide E and protects *C. elegans* against *Bt* infection directly, while MYb115 does not seem to directly inhibit pathogen growth (9). Also, in contrast to MYb115 that only has neutral or beneficial effects on host physiology, MYb11 reduces worm lifespan (31) and aggravates killing upon exposure to purified *Bt* toxins (31). Thus, MYb11 may have a pathogenic potential in some contexts. In line with this thought, we here found that *P. lurida* MYb11 increases the abundance of known pathogen-responsive proteins, while *P. fluorescens* MYb115 does not. These proteins are commonly referred to as *C. elegans* immune defense proteins, albeit the exact function of the majority of these proteins is unknown. We could confirm MYb11-specific activation of expression of the CUB-like domain encoding genes *dod-24* and *F55G11.4* on the transcript level. Interestingly, F55G11.4p::gfp expression is primarily localized to the first intestinal ring (int1). This observation is reminiscent of exclusive expression of some *C. elegans* C-type lectin-like genes such as *clec-42* and *clec-43* in int1 (56). The expression of potential immune effectors specifically by int1 might reflect specialization of int1 as the ‘entry gate’ of the intestine, creating a distinct microenvironment that is important for host-microbe interactions.

The increased abundance of immune effector proteins in the presence of MYb11 indicates that MYb11 activates *C. elegans* pathogen defenses. This may reflect its pathogenic potential but may also contribute to its protective effect against *Bt* infection. Demonstrating the involvement of individual immune effectors in microbiota-mediated protection using knock-outs of single genes can be challenging due to potential functional redundancy or gene compensation among *C. elegans* immune effectors. Indeed, neither mutant of *dod-24*, *lys-1*, or *clec-41* showed reduced protection by MYb11 upon *Bt* exposure. However, several genes encoding the proteins that we found to be modulated by MYb11 are targets of the *C. elegans* p38 MAPK immune and stress signaling pathway (72–75) and recent work by Griem-Krey *et al.* shows that disruption of p38 MAPK signaling not only abolishes, but completely reverses the protective effect of MYb11 upon infection with Bt679 (76). Thus, we hypothesize that in addition to the production of the antimicrobial compound massetolide E that directly inhibits *Bt* growth (9), MYb11 can protect *C. elegans* from pathogen infection by activating its immune defenses.

While we identified a clear MYb11-specific proteome signature that may contribute to its protective effect, identifying MYb115-specific protein targets with a potential role in protection proved more challenging. We found that both, *P. lurida* MYb11 and *P. fluorescens* MYb115, affect *C. elegans* fat metabolism proteins, albeit in different ways. Immune response activation has been repeatedly linked to changes in *C. elegans* fat metabolism. For example, the monounsaturated fatty acid oleate, which is the product of FAT-7 activity, is required for the activation of *C. elegans* pathogen defenses against infection with *E. faecalis*, *Serratia marcescens*, and *P. aeruginosa* (58). Also, the nuclear hormone receptor NHR-49, which is a major regulator of *C. elegans* fat metabolism, mediates *C. elegans* defenses against infection with *E. faecalis* (57), *P. aeruginosa* (77), and *S. aureus* (78). We could show that MYb115 reduces FAT-7 expression. However, our analysis of the *nhr-49(ok2165)* mutant indicates that MYb11-, and MYb115-mediated protection against *Bt* infection is independent of *nhr-49*. Thus, the role of *C. elegans* fat metabolism in microbiota-mediated protection against pathogen infection remains to be determined.

The most interesting candidate proteins that we could identify and that may be involved in MYb115-mediated protection are the intermediate filament proteins of the *C. elegans* cytoskeleton: Several intermediate filaments were more abundant in MYb115-treated worms compared to MYb11- or OP50-exposed worms and the *ifb-2(kc14)* mutant, which lacks an endotube, showed reduced protection by MYb115. In the context of infection, the cytoskeleton functions as a vital barrier against microbial intruders (reviewed in (79, 80)), but can also be modulated by pathogens to support host colonization (reviewed in (81, 82)). We speculate that modulations in cytoskeleton dynamics, i.e., via an increase in intermediate filament protein production, by MYb115 might enhance the integrity of the intestinal barrier and thus contribute to defense against pathogens. Indeed, the *Bt* pore-forming toxin Cry5B leads to structural alterations in the *C. elegans* intermediate filament-rich endotube and the intermediate filament IFB-2 is not only more abundant upon Cry5B exposure, but is also required to withstand the detrimental impact of Cry5B (65). Furthermore, the *C. elegans* NCK-1 homolog to human Nck, an activator of actin assembly, was reported to be required for membrane repair after pore-forming toxin attack (83). Further research is warranted to elucidate the impact of *P. fluorescens* MYb115 on the *C. elegans* intestinal cytoskeleton and its exact role in microbiota-mediated protection against *Bt* pore-forming toxins.

## Acknowledgments

We thank Lena Bluhm, Sabrina Butze, Laura Brügmann, Johanna Jarstorff, and Hanne Griem-Krey for technical support and the Schulenburg group for valuable feedback and discussions. We appreciate the services of the CGC, which is funded by NIH Office of Research Infrastructure Programs (P40 OD010440), for providing worm strains; the Leube Lab at RWTH Aachen, Germany, for providing the mutant *ifb-2(kc14)*, and SunyBiotech for generating transgenic strain F55G11.4p::gfp. This project was funded by the German Science Foundation DFG (Collaborative Research Center CRC 1182 Origin and Function of Metaorganisms, project A1.2 to KD and project Z3 to AT).

## Supplemental Materials

**Figure S1.**
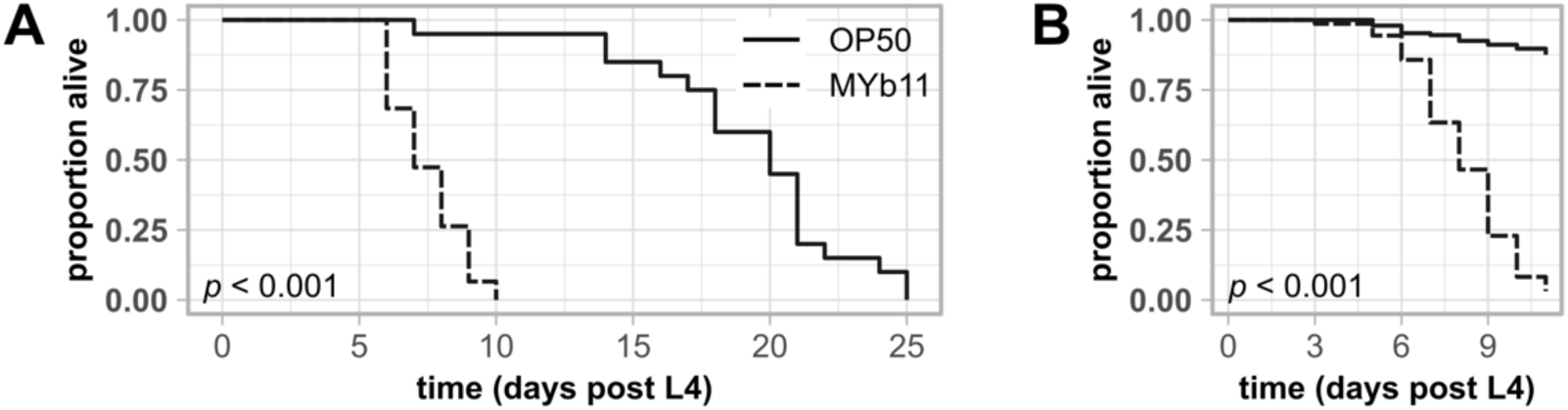
Exposure to *P. lurida* MYb11 reduces the lifespan of wild type N2. Kaplan-Meier curves (84) showing the lifespan of wild type N2 worms on NGM seeded with either OP50 or MYb11. Two independent experiments are shown. Significant differences between worms exposed to OP50 and worms exposed to MYb11 were determined by a log-rank test (85) with (A) individual worms (*n* = 20) and with (B) worm populations of 30 individuals each (*n* = 5).

**Figure S2.**
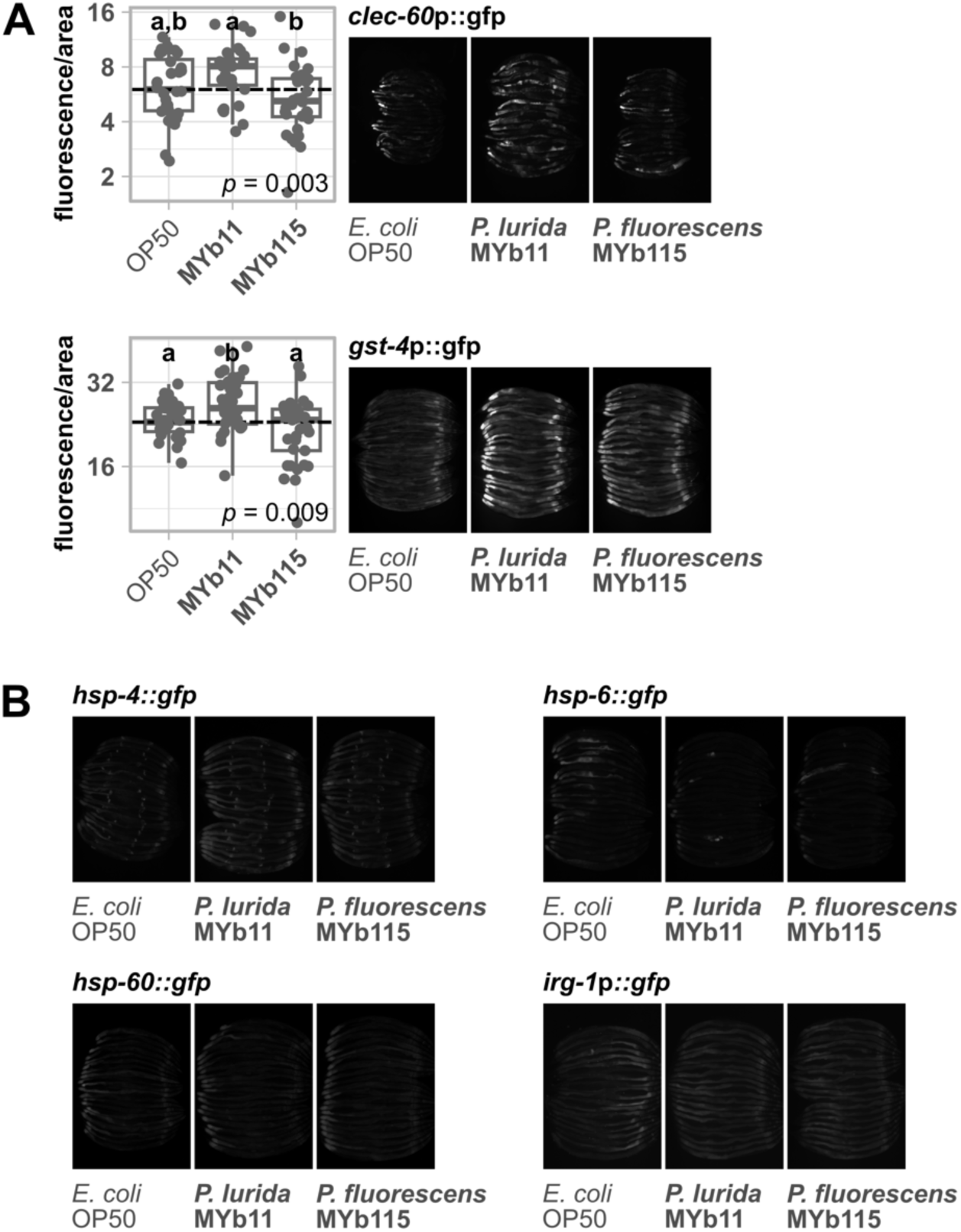
MYb115 does not activate expression of *C. elegans* stress reporters, while MYb11 activates *clec-60*p::gfp and *gst-4*p::gfp expression. Transgenic *C. elegans* reporter strains demonstrating *in vivo* expression of selected genes/promotor sequences tagged with gfp. Transgenic strains were exposed to either *E. coli* OP50, *P. lurida* MYb11, or *P. fluorescens* MYb115, and fluorescent signals imaged in groups of 20 individuals as young adults. Worms were arranged with the heads pointing to the right. (A) The boxplots display the quantification of the gfp fluorescence in young adults (24 h post L4) normalized by the worm’s body size (area). Each dot represents one worm with *n* = 29-35, the dashed line represents the median of the mean grey value for OP50-exposed worms. The *p*-value indicates the statistical significance among the differently exposed worms according to a Kruskal-Wallis rank sum test (32). The *post hoc* Dunn’s test (33) with Bonferroni correction provides the statistical significances between the differently exposed worms and is denoted with letters (same letters indicate no significant differences).

**Figure S3.**
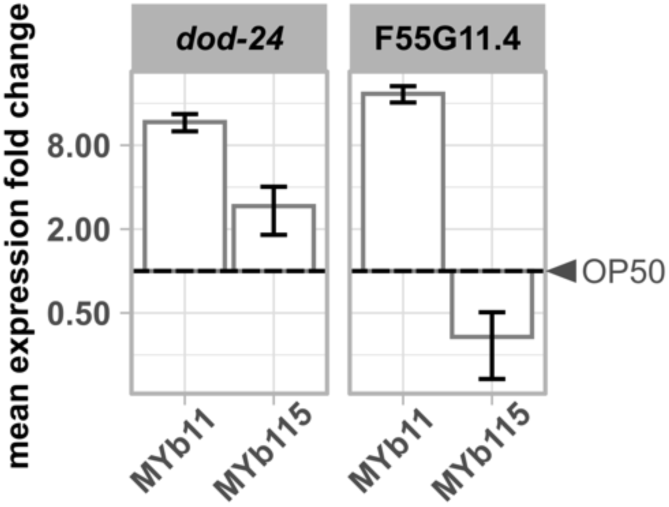
*P. lurida* MYb11 induces the expression of F55G11.4 and *dod-24* more strongly than MYb115. Expression of *dod-24* and F55G11.4 in worms exposed to either MYb11 or MYb115 in relation to worms fed with OP50 (depicted as dashed line) measured with qRT-PCR. Means ± standard deviation (SD) of *n* = 2 are shown. Raw data is provided in Table S6.

**Figure S4.**
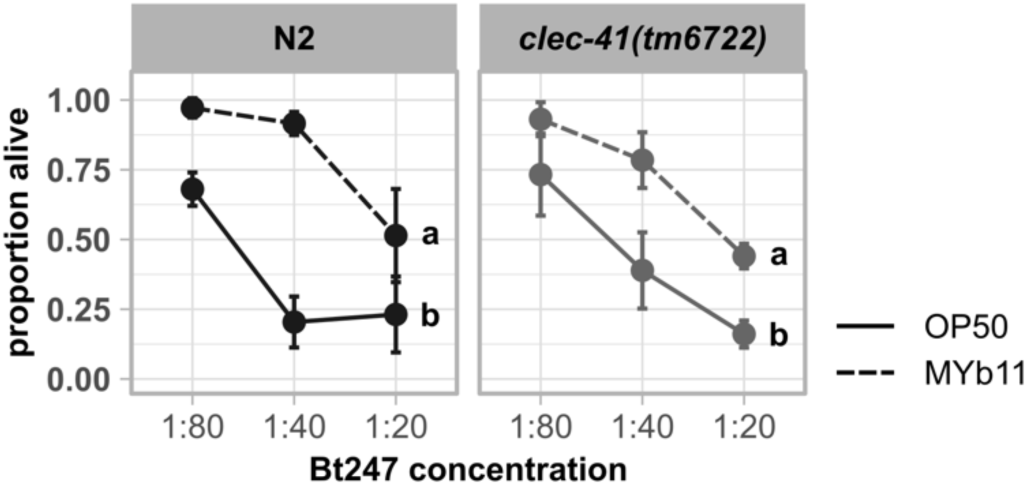
*clec-41* is not required for the protection by MYb11 upon Bt247 exposure. Repetition of experiment in Figure 4D. Survival of mutants *clec-41(tm6722)* and wild type N2 infected with serial dilutions of *B. thuringiensis* Bt247 after 24 hpi (post infection). Worms were exposed to either OP50 or MYb11, before and during infection. Each dot represents the mean ± standard deviation (SD) of four worm populations (*n* = 4). Same letters indicate no significant differences between the dose response curves according to a generalized linear model (GLM) (51) and Bonferroni correction. Raw data and corresponding *p*-values are provided in Table S6.

**Figure S5.**
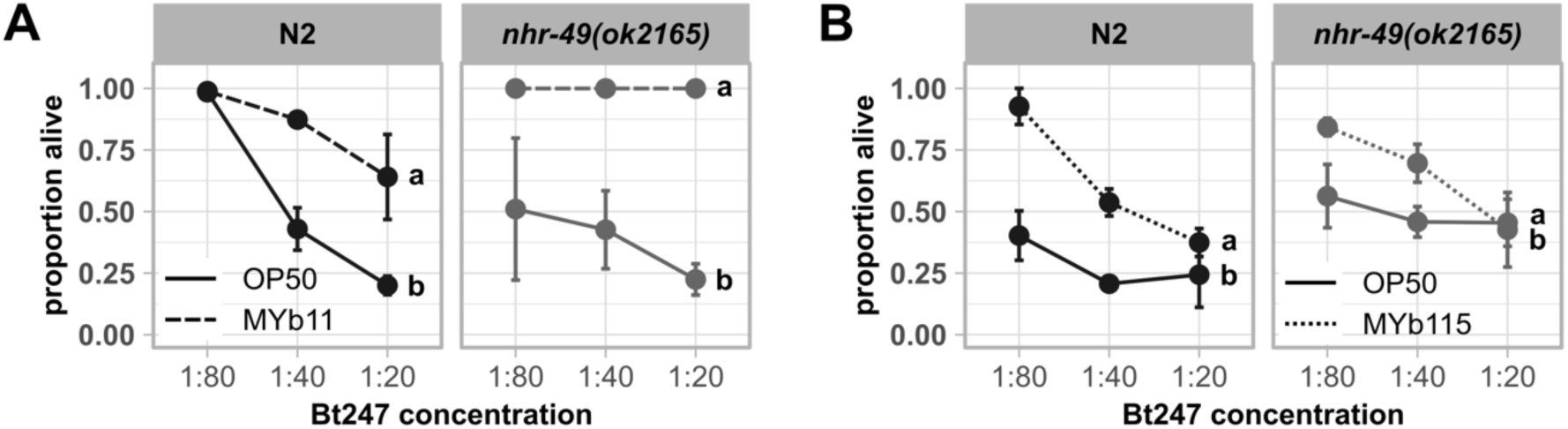
Common fat metabolism regulator NHR-49 is not involved in the defense against Bt247 infection. Repetitions of experiment in Figure 5. Survival of mutant *nhr-49(ok2165)* and wild type N2 infected with serial dilutions of *B. thuringiensis* Bt247 after 24 hpi. Worms were fed with either (A) OP50 and MYb11 or (B) OP50 and MYb115 before and during infection. Each dot represents the mean ± standard deviation (SD) of three worm populations (*n* = 3). Same letters indicate no significant differences between the dose response curves according to a generalized linear model (GLM) (51) and Bonferroni correction. Raw data and corresponding *p*-values are provided in Table S6.

**Figure S6.**
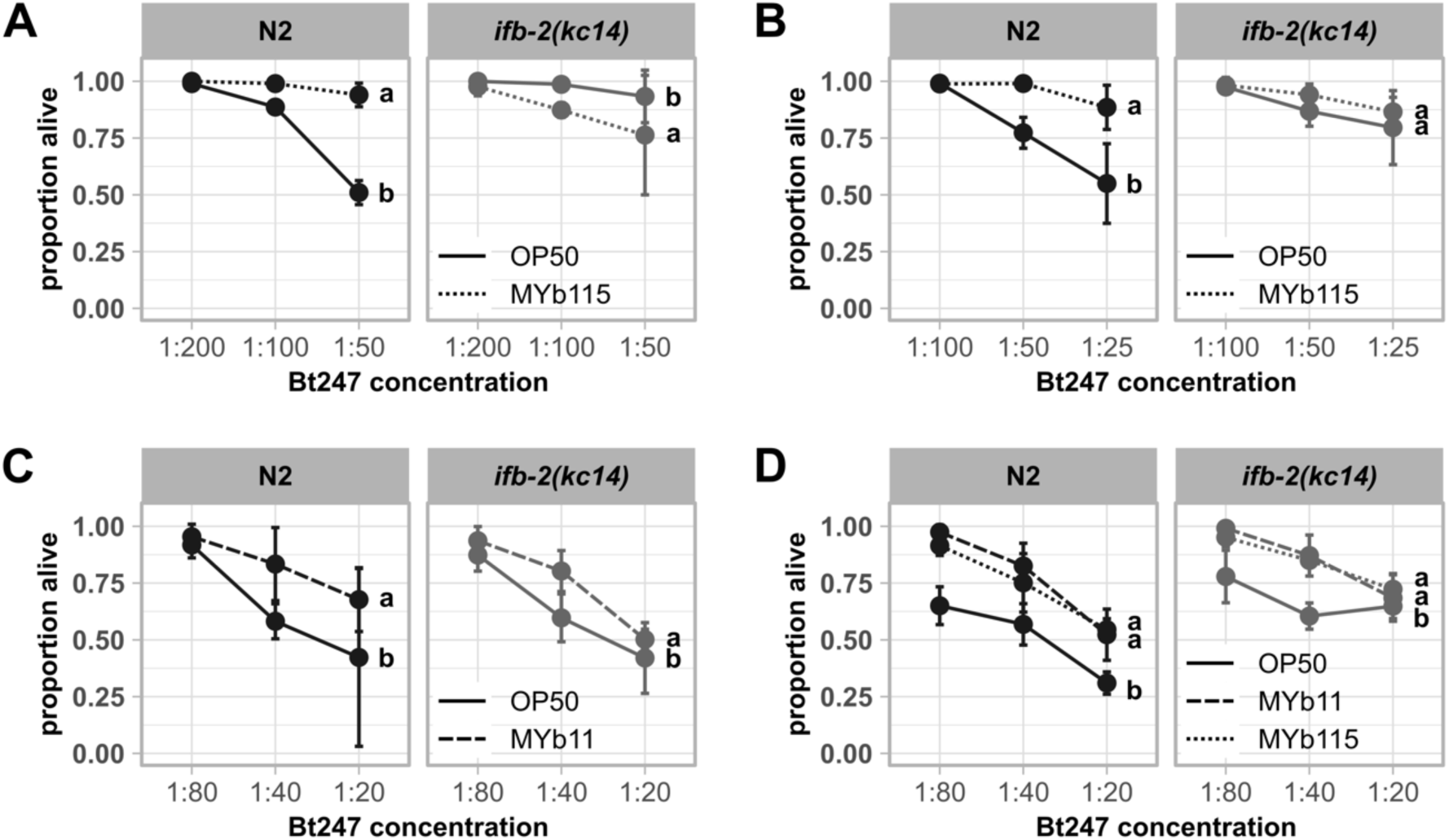
Knock-out of *ifb-2* affects MYb115-mediated protection against Bt247 infection. Repetitions of experiment in Figure 6. Survival of wild type N2 and mutant *ifb-2(kc14)* infected with serial dilutions of *B. thuringiensis* Bt247 after 24 hpi. Worms were exposed to either OP50, MYb11, or MYb115 before and during infection. Each dot represents the mean ± standard deviation (SD) of four worm populations (*n* = 4). Same letters indicate no significant differences between the dose response curves according to a generalized linear model (GLM) (51) and Bonferroni correction. Raw data and corresponding *p*-values are provided in Table S6.

**Table S1. Transgenic strains and mutants employed in this study. Table S2. Primer sequences for qRT–PCRs.**

**Table S3. Raw data and statistical analyses of proteome dataset.**

**Table S4. Significantly enriched gene ontology (GO) terms in cluster 1 and 4 (different abundances of proteins uniquely in MYb11-treated worms).**

**Table S5. Significantly enriched gene ontology (GO) terms in cluster 2 and 3 (different abundances of proteins uniquely in MYb115-treated worms).**

**Table S6. Raw data and statistical analyses of experiments.**

**Supplemental Materials and Methods**

## References

1. Zhang J, Holdorf AD, Walhout AJ. 2017. C. elegans and its bacterial diet as a model for systems-level understanding of host–microbiota interactions. Current Opinion in Biotechnology 46:74–80.

2. Roselli M, Schifano E, Guantario B, Zinno P, Uccelletti D, Devirgiliis C. 2019. *Caenorhabditis elegans* and probiotics interactions from a prolongevity perspective. 20. Int J Mol Sci 20:E5020.

3. Dirksen P, Marsh SA, Braker I, Heitland N, Wagner S, Nakad R, Mader S, Petersen C, Kowallik V, Rosenstiel P, Félix M-A, Schulenburg H. 2016. The native microbiome of the nematode *Caenorhabditis elegans*: gateway to a new host-microbiome model. BMC Biol 14:38.

4. Zhang F, Berg M, Dierking K, Félix M-A, Shapira M, Samuel BS, Schulenburg H. 2017. *Caenorhabditis elegans* as a model for microbiome research. Front Microbiol 8.

5. Dirksen P, Assié A, Zimmermann J, Zhang F, Tietje A-M, Marsh SA, Félix M-A, Shapira M, Kaleta C, Schulenburg H, Samuel B. 2020. CeMbio - The *Caenorhabditis elegans* microbiome resource. G3 g3.401309.2020.

6. Backes C, Martinez-Martinez D, Cabreiro F. 2021. *C. elegans*: A biosensor for host–microbe interactions. Lab Anim 50:127–135.

7. Berg M, Zhou XY, Shapira M. 2016. Host-specific functional significance of *Caenorhabditis* gut commensals. Front Microbiol 7.

8. Montalvo-Katz S, Huang H, Appel MD, Berg M, Shapira M. 2013. Association with soil bacteria enhances p38-dependent infection resistance in *Caenorhabditis elegans*. Infection and Immunity 81:514–520.

9. Kissoyan KAB, Drechsler M, Stange E-L, Zimmermann J, Kaleta C, Bode HB, Dierking K. 2019. Natural *C. elegans* microbiota protects against infection via production of a cyclic lipopeptide of the viscosin group. Current Biology 29:1030–1037.e5.

10. MacNeil LT, Watson E, Arda HE, Zhu LJ, Walhout AJM. 2013. Diet-induced developmental acceleration independent of TOR and insulin in *C. elegans*. Cell 153:240–252.

11. Yang W, Petersen C, Pees B, Zimmermann J, Waschina S, Dirksen P, Rosenstiel P, Tholey A, Leippe M, Dierking K, Kaleta C, Schulenburg H. 2019. The inducible response of the nematode *Caenorhabditis elegans* to members of its natural microbiota across development and adult life. Front Microbiol 10:1793.

12. Coolon JD, Jones KL, Todd TC, Carr BC, Herman MA. 2009. *Caenorhabditis elegans* genomic response to soil bacteria predicts environment-specific genetic effects on life history traits. 6. PLoS Genet 5:e1000503.

13. Goya ME, Xue F, Sampedro-Torres-Quevedo C, Arnaouteli S, Riquelme-Dominguez L, Romanowski A, Brydon J, Ball KL, Stanley-Wall NR, Doitsidou M. 2020. Probiotic *Bacillus subtilis* protects against α-synuclein aggregation in *C. elegans*. Cell Reports 30:367–380.e7.

14. Grishkevich V, Ben-Elazar S, Hashimshony T, Schott DH, Hunter CP, Yanai I. 2012. A genomic bias for genotype-environment interactions in *C. elegans*. Mol Syst Biol 8:587.

15. Cassidy L, Petersen C, Treitz C, Dierking K, Schulenburg H, Leippe M, Tholey A. 2018. The *Caenorhabditis* elegans proteome response to naturally associated microbiome members of the genus *Ochrobactrum*. PROTEOMICS 18:1700426.

16. Mørch MGM, Møller KV, Hesselager MO, Harders RH, Kidmose CL, Buhl T, Fuursted K, Bendixen E, Shen C, Christensen LG, Poulsen CH, Olsen A. 2021. The TGF-β ligand DBL-1 is a key player in a multifaceted probiotic protection against MRSA in C. elegans. Sci Rep 11:10717.

17. Stiernagle T. 2006. Maintenance of *C. elegans*. WormBook 10.1895/wormbook.1.101.1.

18. Borgonie G, Van Driessche R, Leyns F, Arnaut G, De Waele D, Coomans A. 1995. Germination of *Bacillus thuringiensis* spores in bacteriophagous nematodes (Nematoda: Rhabditida). Journal of Invertebrate Pathology 65:61–67.

19. Ou H-L, Kim CS, Uszkoreit S, Wickström SA, Schumacher B. 2019. Somatic niche cells regulate the CEP-1/p53-mediated DNA damage response in primordial germ cells. Developmental Cell 50:167–183.e8.

20. Livak KJ, Schmittgen TD. 2001. Analysis of relative gene expression data using real-time quantitative PCR and the 2^−ΔΔC^ method. Methods 25:402–408.

21. Schneider CA. 2012. NIH Image to ImageJ: 25 years of image analysis. FOCUS ON BIOIMAGE INFORMATICS.

22. Hughes CS, Moggridge S, Müller T, Sorensen PH, Morin GB, Krijgsveld J. 2019. Single-pot, solid-phase-enhanced sample preparation for proteomics experiments. Nat Protoc 14:68–85.

23. Tyanova S, Cox J. 2018. Perseus: A bioinformatics platform for integrative analysis of proteomics data in cancer research, p. 133–148. In von Stechow, L (ed.), Cancer Systems Biology: Methods and Protocols. Springer New York, New York, NY.

24. Vizcaíno JA, Deutsch EW, Wang R, Csordas A, Reisinger F, Ríos D, Dianes JA, Sun Z, Farrah T, Bandeira N, Binz P-A, Xenarios I, Eisenacher M, Mayer G, Gatto L, Campos A, Chalkley RJ, Kraus H-J, Albar JP, Martinez-Bartolomé S, Apweiler R, Omenn GS, Martens L, Jones AR, Hermjakob H. 2014. ProteomeXchange provides globally coordinated proteomics data submission and dissemination. Nat Biotechnol 32:223–226.

25. Cheng X, Yan J, Liu Y, Wang J, Taubert S. 2021. eVITTA: a web-based visualization and inference toolbox for transcriptome analysis. Nucleic Acids Research 49:W207–W215.

26. Wickham H. 2016. ggplot2: Elegant graphics for data analysis. Springer-Verlag New York.

27. Blumenthal T, Squire M, Kirtland S, Cane J, Donegan M, Spieth J, Sharrock W. 1984. Cloning of a yolk protein gene family from *Caenorhabditis elegans*. Journal of Molecular Biology 174:1–18.

28. Spieth J, Blumenthal T. 1985. The *Caenorhabditis elegans* vitellogenin gene family includes a gene encoding a distantly related protein. Molecular and Cellular Biology 5:2495–2501.

29. Perez MF, Lehner B. 2019. Vitellogenins - Yolk gene function and regulation in *Caenorhabditis elegans*. Front Physiol 10:1067.

30. Van Nostrand EL, Sánchez-Blanco A, Wu B, Nguyen A, Kim SK. 2013. Roles of the developmental regulator *unc-62*/Homothorax in limiting longevity in *Caenorhabditis elegans*. PLoS Genet 9:e1003325.

31. Kissoyan KAB, Peters L, Giez C, Michels J, Pees B, Hamerich IK, Schulenburg H, Dierking K. 2022. Exploring effects of *C. elegans* protective natural microbiota on host physiology. Front Cell Infect Microbiol 12:775728.

32. Kruskal WH, Wallis WA. 1952. Use of ranks in one-criterion variance analysis. Journal of the American Statistical Association 47:583–621.

33. Dunn OJ. 1964. Multiple comparisons using rank sums. Technometrics 6:241–252.

34. Zimmermann J, Obeng N, Yang W, Pees B, Petersen C, Waschina S, Kissoyan KA, Aidley J, Hoeppner MP, Bunk B, Spröer C, Leippe M, Dierking K, Kaleta C, Schulenburg H. 2020. The functional repertoire contained within the native microbiota of the model nematode *Caenorhabditis elegans*. ISME J 14:26–38.

35. Watson E, MacNeil LT, Ritter AD, Yilmaz LS, Rosebrock AP, Caudy AA, Walhout AJM. 2014. Interspecies systems biology uncovers metabolites affecting *C. elegans* gene expression and life history traits. Cell 156:759–770.

36. Giese GE, Walker MD, Ponomarova O, Zhang H, Li X, Minevich G, Walhout AJ. 2020. *Caenorhabditis elegans* methionine/S-adenosylmethionine cycle activity is sensed and adjusted by a nuclear hormone receptor. eLife 9:e60259.

37. Bender DA. 2003. Nutritional biochemistry of the vitamins, 2nd ed. Cambridge University Press, Cambridge. https://www.cambridge.org/core/books/nutritional-biochemistry-of-the-vitamins/10B31039A2B0F4B4DC58A89C523FAE97.

38. Bito T, Matsunaga Y, Yabuta Y, Kawano T, Watanabe F. 2013. Vitamin B12 deficiency in *Caenorhabditis elegans* results in loss of fertility, extended life cycle, and reduced lifespan. FEBS Open Bio 3:112–117.

39. Revtovich AV, Lee R, Kirienko NV. 2019. Interplay between mitochondria and diet mediates pathogen and stress resistance in *Caenorhabditis elegans*. PLoS Genet 15:e1008011.

40. Watson E, Olin-Sandoval V, Hoy MJ, Li C-H, Louisse T, Yao V, Mori A, Holdorf AD, Troyanskaya OG, Ralser M, Walhout AJ. 2016. Metabolic network rewiring of propionate flux compensates vitamin B12 deficiency in *C. elegans*. eLife 5:e17670.

41. Bulcha JT, Giese GE, Ali MdZ, Lee Y-U, Walker MD, Holdorf AD, Yilmaz LS, Brewster RC, Walhout AJM. 2019. A persistence detector for metabolic network rewiring in an animal. Cell Reports 26:460–468.e4.

42. Arda HE, Taubert S, MacNeil LT, Conine CC, Tsuda B, Van Gilst M, Sequerra R, Doucette-Stamm L, Yamamoto KR, Walhout AJM. 2010. Functional modularity of nuclear hormone receptors in a *Caenorhabditis elegans* metabolic gene regulatory network. Mol Syst Biol 6:367.

43. Benedetti C, Haynes CM, Yang Y, Harding HP, Ron D. 2006. Ubiquitin-like protein 5 positively regulates chaperone gene expression in the mitochondrial unfolded protein response. Genetics 174:229–239.

44. Haynes CM, Petrova K, Benedetti C, Yang Y, Ron D. 2007. ClpP mediates activation of a mitochondrial unfolded protein response in *C. elegans*. Developmental Cell 13:467–480.

45. Leiers B, Kampkötter A, Grevelding CG, Link CD, Johnson TE, Henkle-Dührsen K. 2003. A stress-responsive glutathione S-transferase confers resistance to oxidative stress in *Caenorhabditis elegans*. Free Radical Biology and Medicine 34:1405–1415.

46. Estes KA, Dunbar TL, Powell JR, Ausubel FM, Troemel ER. 2010. bZIP transcription factor *zip-2* mediates an early response to *Pseudomonas aeruginosa* infection in *Caenorhabditis elegans*. Proc Natl Acad Sci USA 107:2153–2158.

47. Irazoqui JE, Ng A, Xavier RJ, Ausubel FM. 2008. Role for β-catenin and HOX transcription factors in *Caenorhabditis elegans* and mammalian host epithelial-pathogen interactions. Proc Natl Acad Sci USA 105:17469–17474.

48. Samuel BS, Rowedder H, Braendle C, Félix M-A, Ruvkun G. 2016. *Caenorhabditis elegans* responses to bacteria from its natural habitats. Proc Natl Acad Sci USA 113:E3941–E3949.

49. Liu Y, Sellegounder D, Sun J. 2016. Neuronal GPCR OCTR-1 regulates innate immunity by controlling protein synthesis in *Caenorhabditis elegans*. Sci Rep 6:36832.

50. Treitz C, Cassidy L, Höckendorf A, Leippe M, Tholey A. 2015. Quantitative proteome analysis of *Caenorhabditis elegans* upon exposure to nematicidal *Bacillus thuringiensis*. Journal of Proteomics 113:337–350.

51. Nelder JA, Wedderburn RWM. 1972. Generalized linear models. Journal of the Royal Statistical Society Series A (General) 135:370.

52. Bolz DD, Tenor JL, Aballay A. 2010. A conserved PMK-1/p38 MAPK is required in *Caenorhabditis elegans* tissue-specific immune response to Yersinia pestis infection. Journal of Biological Chemistry 285:10832–10840.

53. Mack HID, Kremer J, Albertini E, Mack EKM, Jansen-Dürr P. 2022. Regulation of fatty acid desaturase- and immunity gene-expression by mbk-1/DYRK1A in *Caenorhabditis elegans*. BMC Genomics 23:25.

54. Styer KL, Singh V, Macosko E, Steele SE, Bargmann CI, Aballay A. 2008. Innate immunity in *Caenorhabditis elegans* is regulated by neurons expressing NPR-1/GPCR. Science 322:460–464.

55. Jensen VL, Simonsen KT, Lee Y-H, Park D, Riddle DL. 2010. RNAi screen of DAF-16/FOXO target genes in *C. elegans* links pathogenesis and dauer formation. PLoS ONE 5:e15902.

56. Pees B, Yang W, Kloock A, Petersen C, Peters L, Fan L, Friedrichsen M, Butze S, Zárate-Potes A, Schulenburg H, Dierking K. 2021. Effector and regulator: Diverse functions of *C. elegans* C-type lectin-like domain proteins. PLoS Pathog 17:e1009454.

57. Dasgupta M, Shashikanth M, Gupta A, Sandhu A, De A, Javed S, Singh V. 2020. NHR-49 transcription factor regulates immunometabolic response and survival of *Caenorhabditis elegans* during *Enterococcus faecalis* infection. Infect Immun 88:e00130–20.

58. Anderson SM, Cheesman HK, Peterson ND, Salisbury JE, Soukas AA, Pukkila-Worley R. 2019. The fatty acid oleate is required for innate immune activation and pathogen defense in *Caenorhabditis elegans*. PLoS Pathog 15:e1007893.

59. Watts JL, Browse J. 2000. A palmitoyl-CoA-specific Δ9 fatty acid desaturase from *Caenorhabditis elegans*. Biochemical and Biophysical Research Communications 272:263–269.

60. Gilst MRV, Hadjivassiliou H, Jolly A, Yamamoto KR. 2005. Nuclear hormone receptor NHR-49 controls fat consumption and fatty acid composition in *C. elegans*. PLoS Biol 3:e53.

61. Yang W, Dierking K, Esser D, Tholey A, Leippe M, Rosenstiel P, Schulenburg H. 2015. Overlapping and unique signatures in the proteomic and transcriptomic responses of the nematode *Caenorhabditis elegans* toward pathogenic *Bacillus thuringiensis*. Developmental & Comparative Immunology 51:1–9.

62. Carberry K, Wiesenfahrt T, Windoffer R, Bossinger O, Leube RE. 2009. Intermediate filaments in *Caenorhabditis elegans*. Cell Motil Cytoskeleton 66:852–864.

63. Alberts B, Bray D, Hopkin K, Johnson AD, Lewis J, Raff M, Roberts K, Walter P. 2013. Essential cell biology, 4th ed. Garland Publishing.

64. Coch R, Leube R. 2016. Intermediate filaments and polarization in the intestinal epithelium. Cells 5:32.

65. Geisler F, Coch RA, Richardson C, Goldberg M, Denecke B, Bossinger O, Leube RE. 2019. The intestinal intermediate filament network responds to and protects against microbial insults and toxins. Development dev.169482.

66. Geisler F, Coch RA, Richardson C, Goldberg M, Bevilacqua C, Prevedel R, Leube RE. 2020. Intestinal intermediate filament polypeptides in C. elegans: Common and isotype-specific contributions to intestinal ultrastructure and function. Sci Rep 10:3142.

67. McDonagh A, Crew J, van der Linden AM. 2022. Dietary vitamin B12 regulates chemosensory receptor gene expression via the MEF2 transcription factor in *Caenorhabditis elegans*. 6. G3 Genes|Genomes|Genetics 12:jkac107.

68. Nair T, Chakraborty R, Singh P, Rahman SS, Bhaskar AK, Sengupta S, Mukhopadhyay A. 2022. Adaptive capacity to dietary vitamin B12 levels is maintained by a gene-diet interaction that ensures optimal life span. 1. Aging Cell 21:e13518.

69. Salmela H, Amdam GV, Freitak D. 2015. Transfer of immunity from mother to offspring is mediated via egg-yolk protein vitellogenin. PLoS Pathog 11:e1005015.

70. Fischer M, Regitz C, Kull R, Boll M, Wenzel U. 2013. Vitellogenins increase stress resistance of *Caenorhabditis elegans* after *Photorhabdus luminescens* infection depending on the steroid-signaling pathway. Microbes and Infection 15:569–578.

71. Winter AD, Tjahjono E, Beltrán LJ, Johnstone IL, Bulleid NJ, Page AP. 2022. Dietary-derived vitamin B12 protects *Caenorhabditis elegans* from thiol-reducing agents. BMC Biol 20:228.

72. Alper S, McBride SJ, Lackford B, Freedman JH, Schwartz DA. 2007. Specificity and complexity of the *Caenorhabditis elegans* innate immune response. Molecular and Cellular Biology 27:5544–5553.

73. Troemel ER, Chu SW, Reinke V, Lee SS, Ausubel FM, Kim DH. 2006. p38 MAPK regulates expression of immune response genes and contributes to longevity in *C. elegans*. PLoS Genetics 2:e183.

74. Block DHS, Twumasi-Boateng K, Kang HS, Carlisle JA, Hanganu A, Lai TY-J, Shapira M. 2015. The developmental intestinal regulator ELT-2 controls p38-dependent immune responses in adult *C. elegans*. PLoS Genet 11:e1005265.

75. Pukkila-Worley R, Feinbaum R, Kirienko NV, Larkins-Ford J, Conery AL, Ausubel FM. 2012. Stimulation of host immune defenses by a small molecule protects *C. elegans* from bacterial infection. PLoS Genet 8:e1002733.

76. Griem-Krey H, Petersen C, Hamerich IK, Schulenburg H. 2023. The intricate triangular interaction between protective microbe, pathogen and host determines fitness of the metaorganism. Proc R Soc B 290:20232193.

77. Naim N, Amrit FRG, Ratnappan R, DelBuono N, Loose JA, Ghazi A. 2021. Cell nonautonomous roles of NHR-49 in promoting longevity and innate immunity. Aging Cell 20.

78. Wani KA, Goswamy D, Taubert S, Ratnappan R, Ghazi A, Irazoqui JE. 2021. NHR-49/PPAR-α and HLH-30/TFEB cooperate for *C. elegans* host defense via a flavin-containing monooxygenase. eLife 10:e62775.

79. Geisler F, Leube R. 2016. Epithelial intermediate filaments: guardians against microbial infection? Cells 5:29.

80. Mostowy S, Shenoy AR. 2015. The cytoskeleton in cell-autonomous immunity: structural determinants of host defence. Nat Rev Immunol 15:559–573.

81. Dramsi S, Cossart P. 1998. Intracellular pathogens and the actin cytoskeleton. Annual Review of Cell and Developmental Biology 14:137–66.

82. Bhavsar AP, Guttman JA, Finlay BB. 2007. Manipulation of host-cell pathways by bacterial pathogens. Nature 449:827–834.

83. Sitaram A, Yin Y, Zamaitis T, Zhang B, Aroian RV. 2022. A *Caenorhabditis elegans nck-1* and filamentous actin-regulating protein pathway mediates a key cellular defense against bacterial pore-forming proteins. PLoS Pathog 18:e1010656.

84. Kaplan EL, Meier P. 1958. Nonparametric estimation from incomplete observations. Journal of the American Statistical Association 53:457.

85. Harrington D. 2005. Linear rank tests in survival analysisEncyclopedia of Biostatistics. John Wiley & Sons, Ltd.

