## Supplemental figures and material for "The *C. elegans* proteome response to two protective *Pseudomonas* mutualists"

### Supplemental Material

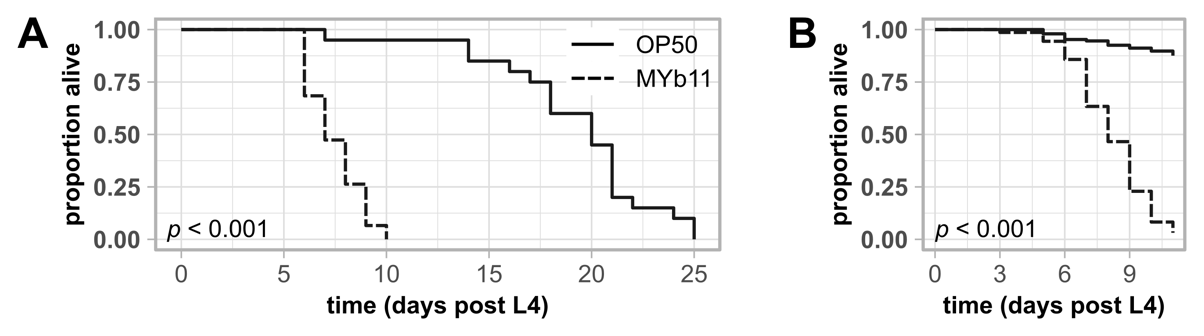

**Figure S1. Exposure to *P. lurida* MYb11 reduces the lifespan of the wild type *C. elegans* N2 strain .** Kaplan-Meier curves (84) showing the lifespan of wild type N2 worms on NGM seeded with either OP50 or MYb11. Two independent experiments are shown. Significant differences between worms exposed to OP50 and worms exposed to MYb11 were determined by a log-rank test (85) with (A) individual worms (*n* = 20) and with (B) worm populations of 30 individuals each (*n* = 5).

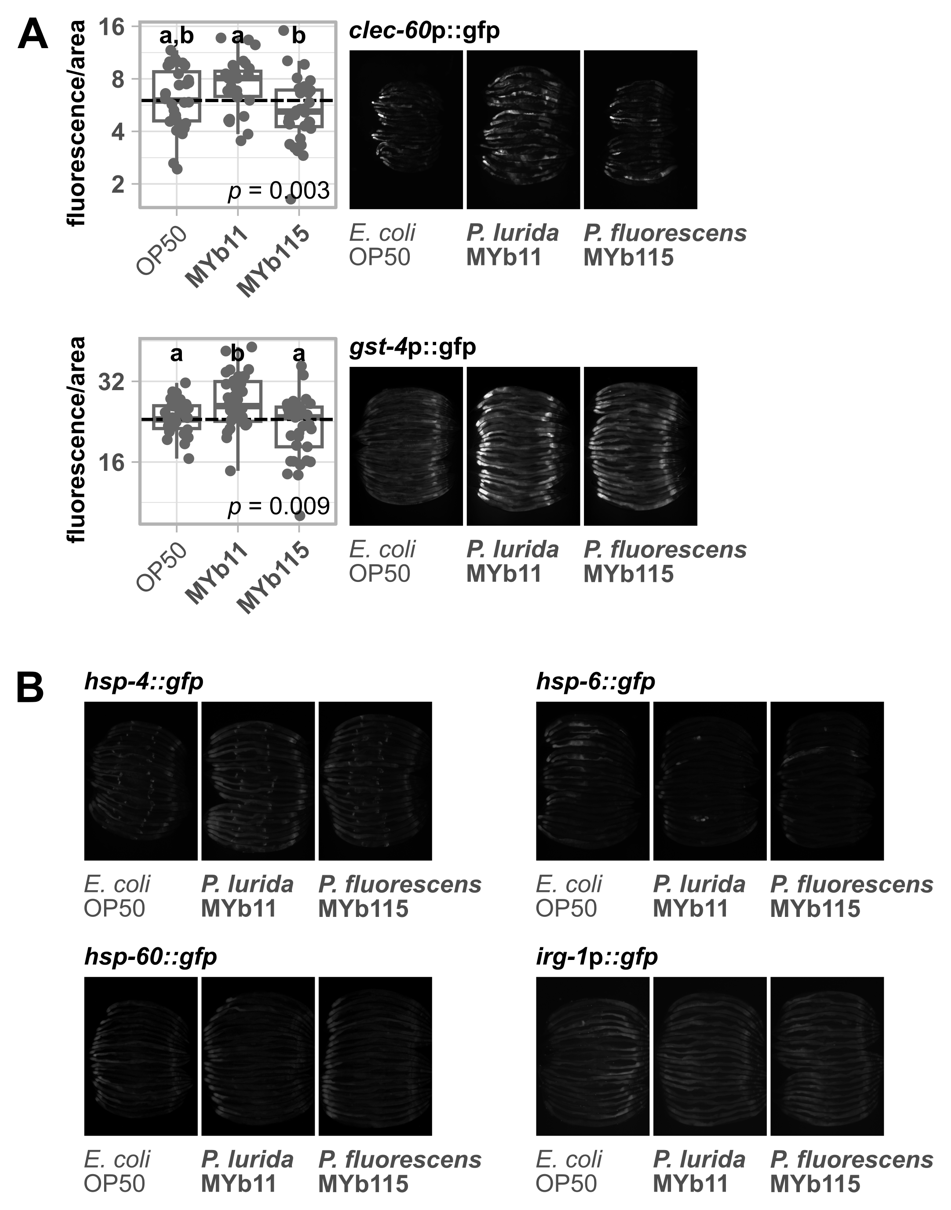

**Figure S2. MYb115 does not activate expression of *C. elegans* stress reporters, while MYb11 activates *clec-60*p::gfp and *gst-4*p::gfp expression.** Transgenic *C. elegans* reporter strains demonstrating *in vivo* expression of selected genes/promotor sequences tagged with gfp. Transgenic strains were exposed to either *E. coli* OP50, *P. lurida* MYb11, or *P. fluorescens* MYb115, and fluorescent signals imaged in groups of 20 individuals as young adults. Worms were arranged with the heads pointing to the right. (A) The boxplots display the quantification of the gfp fluorescence in young adults (24 h post L4) normalized by the worm’s body size (area). Each dot represents one worm with *n* = 29-35, the dashed line represents the median of the mean grey value for OP50-exposed worms. The *p*-value indicates the statistical significance among the differently exposed worms according to a Kruskal-Wallis rank sum test (32). The *post hoc* Dunn’s test (33) with Bonferroni correction provides the statistical significances between the differently exposed worms and is denoted with letters (same letters indicate no significant differences).

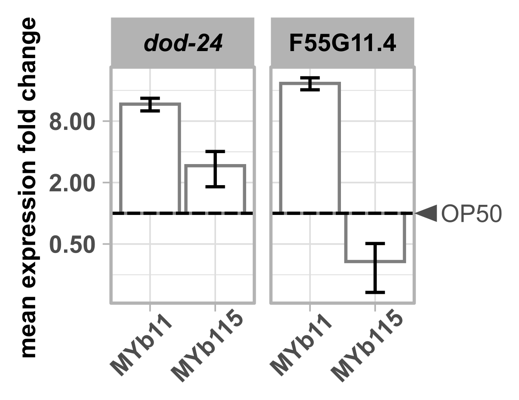

**Figure S3. *P. lurida* MYb11 induces the expression of F55G11.4 and *dod-24* more strongly than MYb115.** Expression of *dod-24* and F55G11.4 in worms exposed to either MYb11 or MYb115 in relation to worms fed with OP50 (depicted as dashed line) measured with qRT-PCR. Means ± standard deviation (SD) of *n* = 2 are shown. Raw data is provided in Table S6.

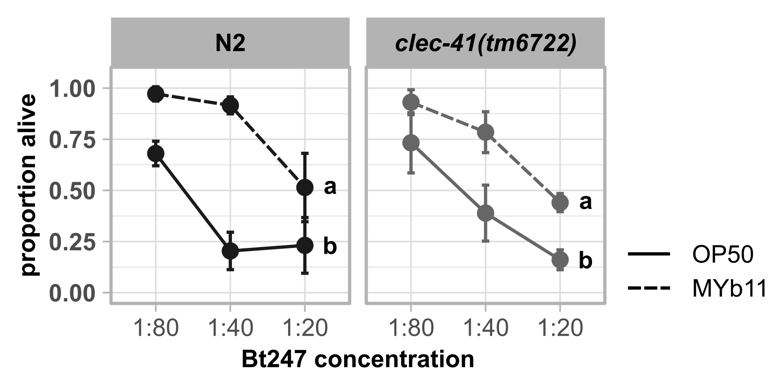

**Figure S4. *clec-41* is not required for the protection by MYb11 upon Bt247 exposure.** Repetition of experiment in Figure 4D. Survival of mutants *clec-41(tm6722)* and wild type N2 infected with serial dilutions of *B. thuringiensis* Bt247 after 24 hpi (post infection). Worms were exposed to either OP50 or MYb11, before and during infection. Each dot represents the mean ± standard deviation (SD) of four worm populations (*n* = 4). Same letters indicate no significant differences between the dose response curves according to a generalized linear model (GLM) (51) and Bonferroni correction. Raw data and corresponding *p*-values are provided in Table S6.

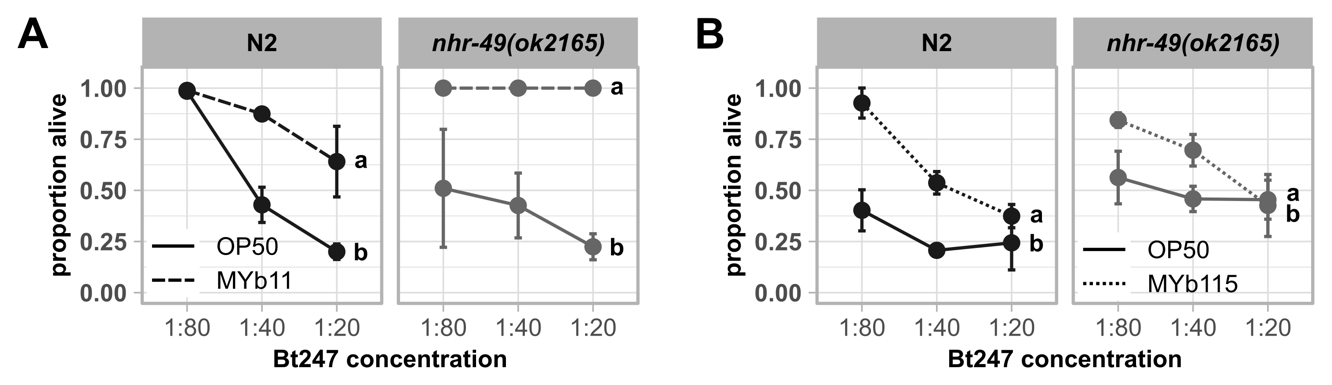

**Figure S5. Common fat metabolism regulator NHR-49 is not involved in the defense against Bt247 infection.** Repetitions of experiment in Figure 5. Survival of mutant *nhr-49(ok2165)* and wild type N2 infected with serial dilutions of *B. thuringiensis* Bt247 after 24 hpi. Worms were fed with either (A) OP50 and MYb11 or (B) OP50 and MYb115 before and during infection. Each dot represents the mean ± standard deviation (SD) of three worm populations (*n* = 3). Same letters indicate no significant differences between the dose response curves according to a generalized linear model (GLM) (51) and Bonferroni correction. Raw data and corresponding *p*-values are provided in Table S6.

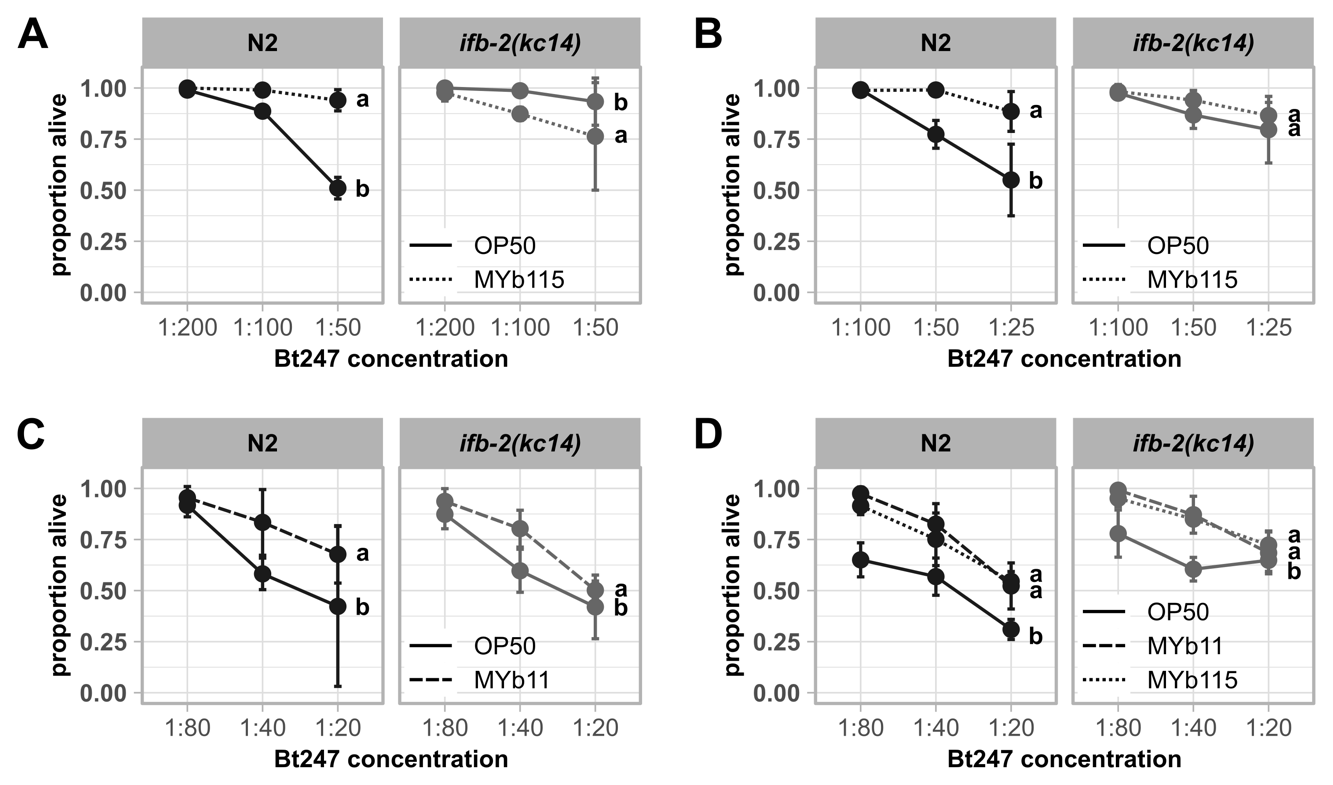

**Figure S6. Knock-out of *ifb-2* affects MYb115-mediated protection against Bt247 infection .** Repetitions of experiment in Figure 6. Survival of wild type N2 and mutant *ifb-2(kc14)* infected with serial dilutions of *B. thuringiensis* Bt247 after 24 hpi. Worms were exposed to either OP50, MYb11, or MYb115 before and during infection. Each dot represents the mean ± standard deviation (SD) of four worm populations (*n* = 4). Same letters indicate no significant differences between the dose response curves according to a generalized linear model (GLM) (51) and Bonferroni correction. Raw data and corresponding *p*-values are provided in Table S6.

### Supplemental Materials & Methods

**LC-MS analysis**

Digested proteomes were analyzed by LC-MS/MS. A Dionex U3000 uHPLC system was coupled to an Orbitrap Fusion Lumos mass spectrometer (Thermo Fisher Scientific). The digested samples were injected on a C18 PepMap 100 precolumn (column dimensions: 300 μm i. d. x 5 mm; Thermo Scientific) with a flow rate of 30 μL/min, trapped and desalted for 2 min and then separated on an Acclaim PepMap RSLC column (column dimension: 75 μm i. d. x 50 cm; Thermo Scientific) over a gradient of eluent A (0.05% aqueous FA) and eluent B (80% ACN, 0.04% FA) with a flow rate of 0.3 μL/min.

Approximately 1 µg of peptide were eluted over a gradient from 5% eluent B to 27% eluent B in 125 min, then to 40% B in 35 min, followed by an increase to 95% eluent B in 20 min. After isocratic elution at 95% eluent B for 18 min, the column was equilibrated for 10 min with 5% eluent B. After each sample LC run, the column was washed using a blank run, injecting 5 μL of loading buffer.

Full MS scans were acquired from 2 min to 200 min in positive ion mode with a resolution of 120,000, the AGC target was set to 4e5, the maximum injection time was 50 ms and the scan range was from 400 m/z to 1100 m/z. Data dependent FT-MS/MS spectra of the most intense precursor ions were acquired for 3 s with a resolution of 7,500; The isolation window was set to 1.2 m/z, the normalized collision energy was 30, the AGC target was 1e5 and a maximum injection time of 50 ms. Precursors with a charge state of 2- 7 were accepted and isotopes were excluded. Dynamic exclusion was set to 60 s with a mass tolerance of 10 ppm. Apex detection properties were set to an expected peak width (TWHM) of 25 s and a desired apex window of 35%.

With three biological conditions, each in 4 replicates, overall 12 samples were analyzed by 1D/LC MS.

**Database search**

The spectral data was searched against a combined database downloaded from UniProt (23.11.2022), which included all the proteins of the different bacteria used on culture plates and those from *C. elegans*. The reference proteome of *C. elegans* (UP000001940; 26,738 entries) were combined with the UniParc entries of *P. lurida* (UP000238409; 5392 entries), *P. fluorescens* (UP000694054; 5548 entries) and *E. coli* OP50 (UP000295638; 4227 entries).

MS raw files were queried against a protein database of target organisms and common contaminants using the Sequest search algorithm and the Proteome discoverer software v. 2.5 (Thermo Scientific). The spectra were recalibrated by a survey search with 20 ppm precursor mass tolerance and 0.5 Da fragment mass tolerance. Recalibration of retention times was performed by non-linear regression; the parameter tuning for the regression was set to fine. Searches with a full-tryptic protease specificity and allowing for variable oxidation of methionine and fixed carbamidomethylation of cysteine residues were rescored using INFERYS and restricted to a false discovery rate below 1% for peptide spectrum matches (PSM) using the Percolator q-value. The target FDR of both peptide group and protein group identifications was set to 1%.

Label-free quantification was performed using the Minora Feature Detector with the following parameters; the minimum trace length was set to 4, and the maximum Δ RT of isotope pattern multiplets was set to 0.2 min. Unique and razor peptides were used for quantification. The precursor abundance was based on the peak area. The protein abundance values were median normalized and log_2_ transformed for statistical analysis.

Prior to the statistical evaluation and hypothesis testing the dataset was filtered to protein group identifications quantified in all four replicates of at least one of the growth conditions. The statistical analysis was performed with and without data imputation, respectively. For data imputation missing values were replaced by low abundance resampling from a normal distribution (width = 0.3; down-shift = 1.8).
